# A terminal selector prevents a Hox transcriptional switch to safeguard motor neuron identity throughout life

**DOI:** 10.1101/643320

**Authors:** Weidong Feng, Yinan Li, Pauline Dao, Jihad Aburas, Priota Islam, Benayahu Elbaz, Anna Kolarzyk, André E.X. Brown, Paschalis Kratsios

## Abstract

Nervous system function critically relies on continuous expression of neuron type-specific terminal identity features, such as neurotransmitter receptors, ion channels and neuropeptides. How individual neuron types select such features during development and maintain them throughout life is poorly understood. Here, we report an unconventional mechanism that enables cholinergic motor neurons (MNs) in the *C. elegans* ventral nerve cord to select and maintain their distinct terminal identity features. The conserved terminal selector UNC-3 (Collier/Ebf) UNC-3 is continuously required not only to promote cholinergic MN features, but also to prevent expression of “unwanted” terminal identity features normally reserved for other neuron types. Mechanistically, this dual function is achieved by the ability of UNC-3 to prevent a switch in the transcriptional targets of the Hox protein LIN-39 (Scr/Dfd/Hox4-5). The strategy of a terminal selector preventing a Hox transcriptional switch may constitute a general principle for safeguarding neuronal terminal identity features throughout life.

## INTRODUCTION

Precise establishment and maintenance of neuron type-specific gene expression programs is essential for nervous system function. Integral components of these programs are effector genes encoding proteins critical for neuronal function (e.g., neurotransmitter [NT] biosynthesis components, ion channels, NT receptors, neuropeptides) [1–4]. Such effector genes, referred to as terminal identity genes herein, are expressed continuously, from development throughout life, in post-mitotic neurons in a combinatorial fashion [2]. Hence, it is the unique overlap of many effector gene products in a specific neuron type that determines each neuron’s distinct terminal identity, and thereby function. But how do post-mitotic neurons select which terminal identity genes to express and which ones to repress? Understanding how neuron type-specific batteries of terminal identity genes are established during development and, perhaps most importantly, maintained throughout life represents one key step towards understanding how individual neuron types become and remain functional. Providing molecular insights into this fundamental problem may also have important biomedical implications, as defects in terminal identity gene expression are associated with a variety of neurodevelopmental and neurodegenerative disorders [1, 5–7].

Seminal genetic studies in multiple model systems revealed a widely employed principle: neuron type-specific transcription factors (TFs) coordinate the expression of “desired” terminal identity genes with the exclusion of “unwanted” terminal identity genes [8–15]. As such, these TFs exert a dual role, as they are not only required to induce a specific set of terminal identity features critical for the function of a given neuron type, but also to simultaneously prevent expression of terminal features normally reserved for other neuron types. Consequently, neurons lacking these TFs fail to acquire their unique terminal identity, and concomitantly gain features indicative of alternative identities. For example, mouse striatal cholinergic interneurons lacking *Lhx7* lose their terminal identity and acquire molecular features indicative of GABAergic interneuron identity [13]. In the midbrain, removal of *Gata2* results in loss of GABAergic neuron identity and simultaneous gain of terminal identity features specific to glutamatergic neurons [12].

However, the molecular mechanisms underlying the dual function of neuron type-specific TFs remain poorly defined. How can the same TF, within the same cell, promote a specific identity and simultaneously prevent “unwanted” terminal features? In principle, the same TF can simultaneously operate as direct activator of neuron type-specific terminal identity genes and direct repressor of alternative identity genes, as exemplified by the case of *Fezf2*, a TF necessary for neocortical projection neuron identity [16]. Another possibility is indirect regulation. For example, neuron type-specific TFs can prevent adoption of alternative identities by repressing expression of intermediary TFs. Supporting this scenario, Tlx1 and Tlx3 jointly promote glutamatergic identity features in dorsal spinal cord neurons and simultaneously repress Pax2, a TF required for terminal identity of GABAergic neurons [11]. However, it remains unclear whether these two mechanisms (direct and indirect regulation) underlying the function of neuron type-specific TFs are broadly applicable in the nervous system.

Although the aforementioned studies begin to explain how neurons select their terminal identity features during development[8–15], the function of neuron type-specific TFs is rarely assessed during post-natal stages. As such, the molecular mechanisms that maintain expression of neuronal terminal identity features are largely unknown. Is the same neuron type-specific TF continuously required, from development through adulthood, to induce a specific set of terminal identity genes and simultaneously prevent “unwanted” features? Alternatively, a given neuron type can employ different mechanisms for selection (during development) and maintenance (through adulthood) of terminal identity features. Addressing this fundamental problem has been challenging in the vertebrate nervous system, in part due to its inherent complexity and difficulty to track individual neuron types with single-cell resolution from embryonic development to adulthood.

To study how neurons select and maintain their terminal features, we use as a model the well-defined motor neuron (MN) subtypes of the *Caenorhabditis elegans* ventral nerve cord (equivalent to spinal cord). Five cholinergic (DA, DB, VA, VB, AS) and two GABAergic (DD, VD) MN subtypes are located along the nerve cord and control locomotion (**Fig. 1A**) [17, 18]. Because they are present in both *C. elegans* sexes (males and hermaphrodites), we will refer to them as “sex-shared” MNs. In addition, there are two subtypes of “sex-specific” cholinergic MNs; the hermaphrodite-specific VC neurons control egg laying [19, 20], whereas the male-specific CA neurons are required for mating [21] (**Fig. 1A**). Apart from distinct morphology and connectivity, each subtype can be molecularly defined based on the combinatorial expression of known terminal identity genes, such as ion channels, NT receptors, and neuropeptides (**Fig. 1B**). An extensive collection of transgenic reporter animals for MN subtype-specific terminal identity genes is available, thereby providing a unique opportunity to investigate, at single-cell resolution, the effects of TF gene removal on developing and adult MNs.

**Figure 1:**
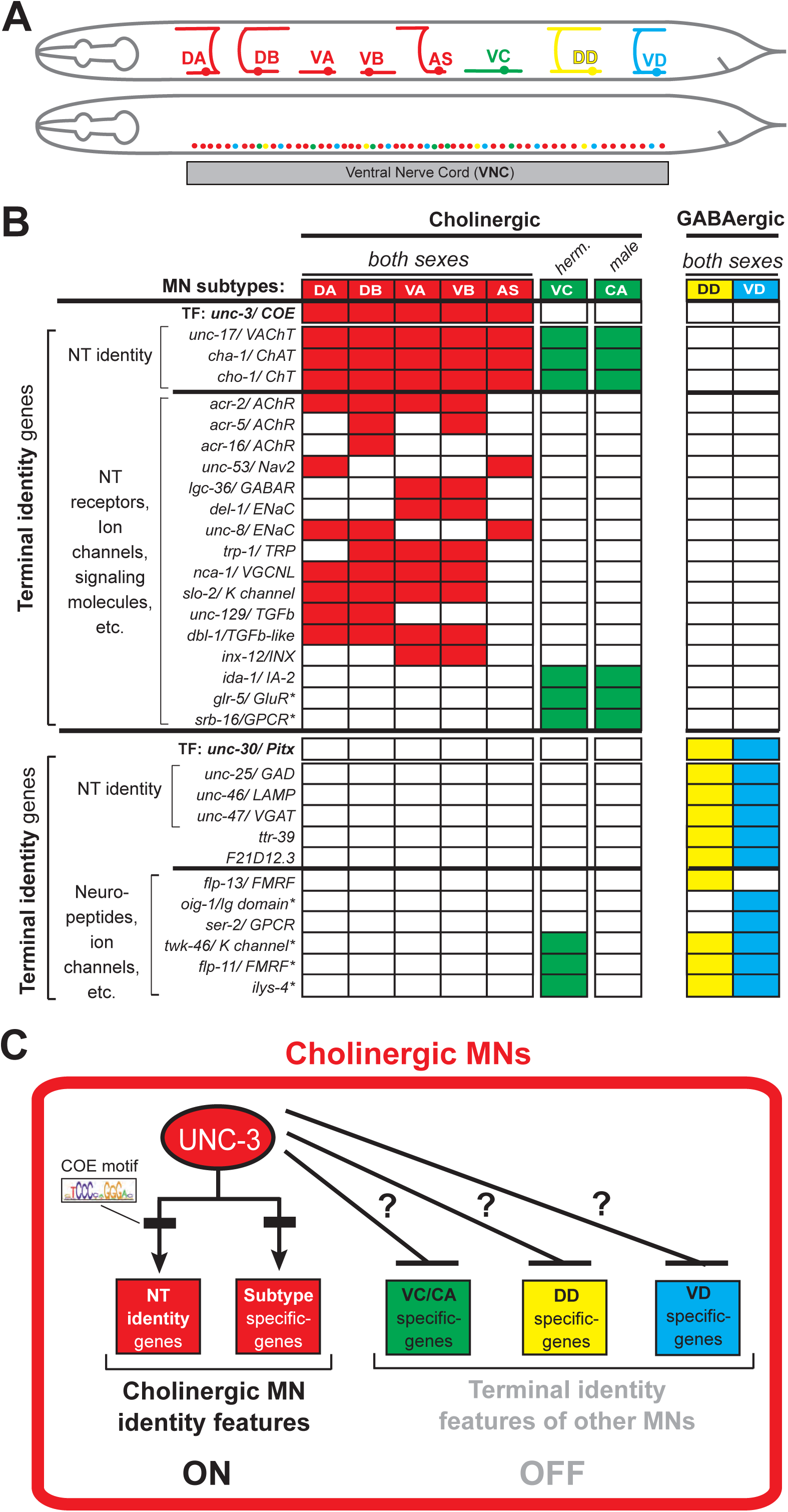
An extensive collection of terminal identity markers for distinct motor neuron subtypes of the *C. elegans* ventral nerve cord. **A:** Schematic showing distinct morphology for each motor neuron subtype in the *C. elegans* hermaphrodite. Below, colored dots represent the invariant cell body position of all MNs of the ventral nerve cord (VNC). Red: 39 sex-shared cholinergic MNs (DA2-7 = 6 neurons, DB3-7 = 5, VA2-11 = 10, VB3-11 = 9, AS2-10 = 9); Green: 6 hermaphrodite-specific VC MNs; Yellow: 4 sex-shared GABAergic DD neurons (DD2-5 = 4); Blue: 9 sex-shared GABAergic VD neurons (VD3-11 = 9). With the exception of VC, all other subtypes have 1-3 extra neurons located at the flanking ganglia (retrovesicular and pre-anal) of the VNC (not shown). Individual neurons of each subtype intermingle along the VNC. **B:** Table summarizing expression of terminal identity markers for VNC MNs. The sex-shared GABAergic MNs (DD, VD) and the sex-specific MNs (VC, CA) do not express UNC-3. Conversely, the sex-shared cholinergic MNs (DA, DB, VA, VB, AS) and the sex-specific MNs (VC, CA) do not express UNC-30/Pitx. For the genes indicated with an asterisk (*), a detailed expression pattern is provided in **Suppl. Fig. 1**. Of note, the male-specific MNs of the CP subtype are also not shown. **C:** Schematic that summarizes the known function of UNC-3 (activator of cholinergic MN identity genes) and the question under investigation: does UNC-3 prevent expression of terminal identity features reserved for other MN subtypes?

UNC-3, the sole *C. elegans* ortholog of the Collier/Olf/Ebf (COE) family of TFs, is selectively expressed in all sex-shared cholinergic MNs of the nerve cord (**Fig. 1B**)[22–26]. Animals lacking *unc-3* display striking locomotion defects [27]. UNC-3 is known to directly activate a large battery of terminal identity genes expressed either in all sex-shared cholinergic MNs (e.g., the NT identity genes *unc-17/* VAChT and *cha-1/* ChAT), or in certain subtypes (e.g., ion channels, NT receptors, signaling molecules) [23] (**Fig. 1B-C**). Based on its ability to broadly co-regulate many distinct terminal identity features, *unc-3* has been classified as a terminal selector gene [2]. Besides its well-established function as activator of terminal identity genes in cholinergic MNs, whether and how UNC-3 can prevent expression of “unwanted” terminal features remains unclear.

Here, we identify a dual role for UNC-3 and describe an unconventional mechanism through which sex-shared cholinergic MNs select and maintain their terminal identity features. We find that UNC-3 is continuously required, from development through adulthood, not only to activate cholinergic MN identity genes, but also to prevent expression of terminal features normally reserved for three other neuron types of the nerve cord (VD, VC, CA). However, MNs lacking *unc-3* do not adopt a mixed identity, i.e., a composite of ectopically expressed features of VD, VC, and CA neurons. Instead, our single-cell analysis of an extensive repertoire of MN terminal identity markers revealed two distinct populations of *unc-3*-depleted MNs. One population gains terminal features of sex-shared GABAergic VD neurons, while a second population acquires terminal features of sex-specific MNs, i.e., CA features in *unc-3* males and VC features in *unc-3* hermaphrodites. Through an unbiased genetic screen, we identified the Hox protein LIN-39 (Scr/Dfd/Hox4-5) as the intermediary factor necessary for induction and maintenance of both sex-shared (VD) and sex-specific terminal identity features in *unc-3* mutants. Intriguingly, suppression of “unwanted” identity features (e.g., VD, VC) relies on a mechanism that does not involve repression of the intermediary factor LIN-39/Hox. Removal of *unc-3* enables LIN-39 to activate “unwanted” terminal identity features instead of cholinergic MN identity genes. Hence, UNC-3 prevents a switch in the transcriptional targets of LIN-39/Hox in cholinergic MNs. Given that terminal selectors and Hox proteins are known to be expressed in a multitude of neuron types across species [1, 28–30], the strategy of a terminal selector preventing a Hox transcriptional switch may constitute a general principle for safeguarding neuronal terminal identity throughout life.

## RESULTS

### UNC-3 has a dual role in distinct populations of ventral nerve cord (VNC) motor neurons

Neuron type-specific TFs often promote a specific identity and simultaneously suppress features reserved for other, functionally related neuronal types [31]. To test whether UNC-3 prevents expression of features normally reserved for other MN subtypes (**Fig. 1C**), it was essential to identify a set of terminal identity markers for all *unc-3*-negative MN subtypes of the VNC, namely the GABAergic (VD, DD) and sex-specific (VC, CA) MNs (**Fig. 1B**). We undertook a candidate gene approach and examined the precise expression pattern of terminal identity genes (e.g., NT receptors, signaling proteins, ion channels, neuropeptides) reported to be expressed in *unc-3*-negative MNs (www.wormbase.org). We carefully characterized at single-cell resolution the expression of 15 terminal identity genes in wild-type animals of both *C. elegans* sexes at the fourth larval stage (L4) (see Methods and **Suppl. Fig. 1**). This analysis revealed 9 markers highly specific for *unc-3*-negative MNs that fall into four categories (**Fig. 1B**): (**i**) two VD-specific terminal identity markers (*ser-2* / serotonin receptor [ortholog of HTR1D]*; oig-1 /* one Ig domain protein), [32] one DD-specific terminal identity marker (*flp-13 /* FMRF-like neuropeptide), (**iii**) three markers for sex-specific (VC in hermaphrodites, CA in males) MNs (*glr-5* / glutamate receptor [ortholog of GRID/GRIK]; *srb-16* / serpentine GPCR receptor; *ida-1* / ortholog of protein tyrosine phosphatase PTPRN), and (**iv**) three markers expressed in both GABAergic subtypes (DD, VD) and sex-specific MNs (*flp-11* / FRMR-like neuropeptide*, twk-46* / potassium channel [ortholog of KCNK1]*, ilys-4 /* invertebrate-type lysozyme).

These 9 markers enabled us to test whether *unc-3-*depleted MNs gain expression of terminal features normally reserved for other MN subtypes. By using animals carrying a strong loss-of-function (null) allele for *unc-3 (n3435)* [25], we first assessed any putative effects on terminal markers for the sex-shared GABAergic MNs (DD, VD). Although the DD-specific marker *flp-13* is unaffected (**Suppl. Fig. 2A**), ectopic expression of the VD-specific terminal identity markers (*ser-2, oig-1*) was observed in *unc-3*-depleted MNs (**Fig. 2A-B**). Interestingly, this ectopic expression was region-specific, observed only in cholinergic MNs at the mid-body region of the VNC. Importantly, 12.1 ± 2.6 (mean ± STDV) out of the 39 *unc-3*-depleted MNs in the VNC were ectopically expressing these VD markers, suggesting that not all *unc-3-*depleted MNs acquire VD terminal identity features. Given that GABAergic and cholinergic MNs are generated in normal numbers in *unc-3* animals [23], the increase in the number of neurons expressing the VD markers cannot be attributed to early developmental defects affecting MN numbers. We next asked whether these ∼12 MNs adopt additional VD terminal identity features, such as expression of genes involved in GABA biosynthesis (*unc-25/GAD* and *unc-47/VGAT)*, or selectively expressed in GABAergic MNs (*ttr-39, klp-4*). However, this does not appear to be the case (**Suppl. Fig. 2A**). We conclude that, in the absence of *unc-3,* cholinergic MNs not only lose their original terminal identity, but a third of them (∼12 out of 39) in the mid-body VNC region also gain some terminal identity features normally reserved for the sex-shared VD neurons (**Fig. 1**). We will refer to these *unc-3*-depleted MNs as “VD-like” (**Fig. 2G**).

**Figure 2:**
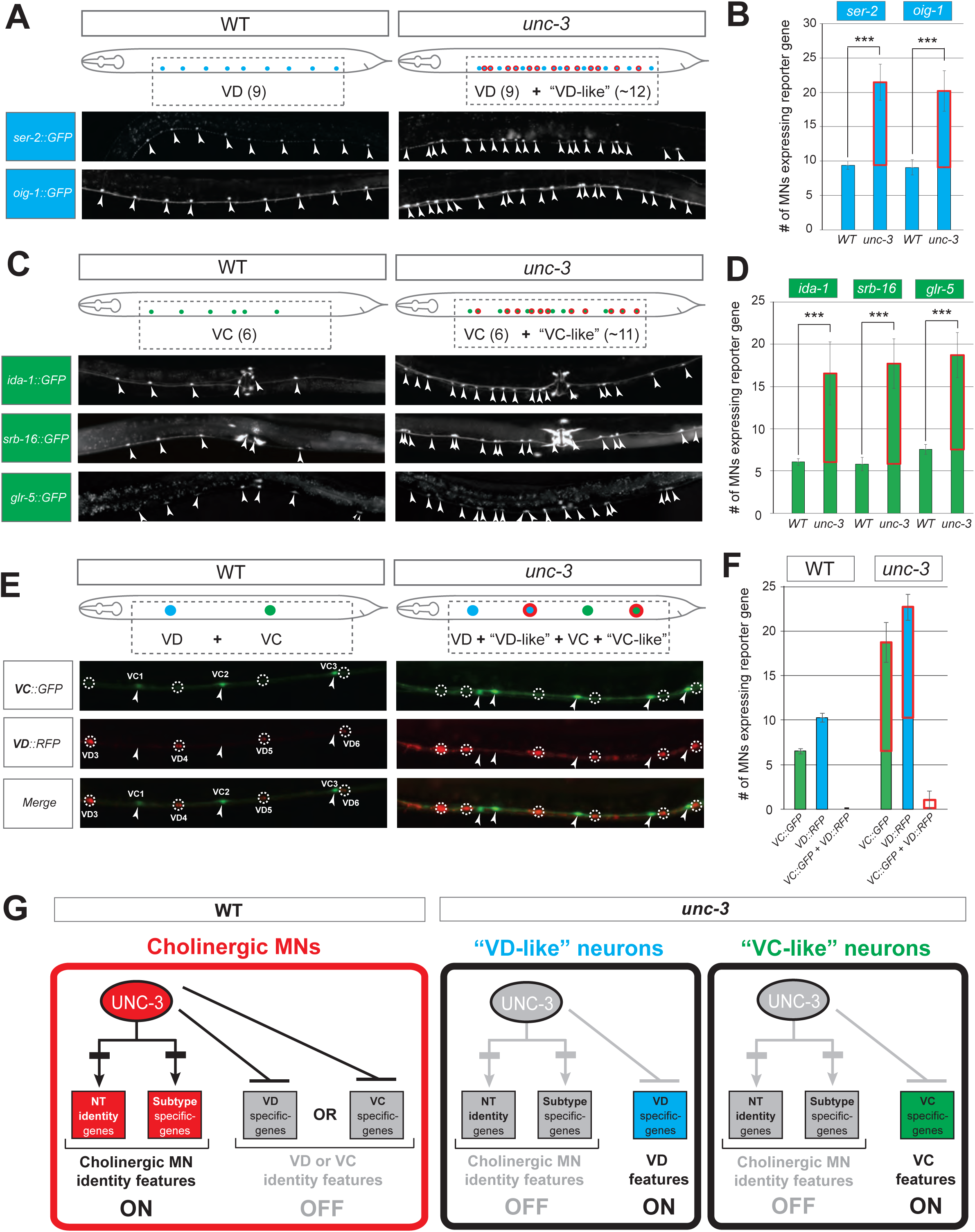
UNC-3 has a dual role in cholinergic ventral cord motor neurons. **A:** Terminal identity markers of VD neurons (*ser-2, oig-1*) are ectopically expressed in *unc-3*-depleted cholinergic MNs. Representative images of larval stage 4 (L4) hermaphrodites are shown. Similar results were obtained in adult animals. Arrowheads point to MN cell bodies with *gfp* marker expression. Green fluorescence signal is shown in white for better contrast. Dotted black box indicates imaged area. **B**: Quantification of VD markers (*ser-2, oig-1*) in WT and *unc-3 (n3435)* at L4. Ectopic expression is highlighted with a red rectangle. N > 15. *** p < 0.001. **C:** Terminal identity markers of VC neurons (*ida-1, srb-16, glr-5*) are ectopically expressed in *unc-3*-depleted cholinergic MNs. Representative images of larval stage 4 (L4) hermaphrodites are shown. Similar results were obtained in adult animals. Arrowheads point to MN cell bodies with *gfp* marker expression. Green fluorescence signal is shown in white for better contrast. Dotted black box indicates imaged area. **D**: Quantification of VC markers (*ida-1, srb-16, glr-5*) in WT and *unc-3 (n3435)* at L4. Ectopic expression in *unc-3*-depleted MNs is highlighted with a red rectangle. N > 15. *** p < 0.001. **E:** Distinct MNs acquire VC-like and VD-like terminal identity features in *unc-3 (n3435).* The VC marker in green (*ida-1::gfp*) and the VD marker in red (*ser-2::rfp*) do not co-localize in WT and *unc-3 (n3435)* mutants. Representative images are shown. Individual VC/VC-like and VD/VD-like neurons are circled (VC: dotted circles; VD: arrowheads) to highlight that an individual MN never expresses both markers. **F:** Quantification of data shown in E. N > 16. **G:** Schematic that summarizes the dual role of *unc-3*. Apart from activating cholinergic MN terminal identity genes, UNC-3 prevents expression of VD and VC terminal features in distinct cells (“VD-like” versus “VC-like”).

To test whether UNC-3 also prevents expression of terminal identity features of sex-specific cholinergic MNs, we examined three VC-specific terminal markers (*glr-5, srb-16, ida-1* in **Fig. 1B**) in hermaphrodite nematodes lacking *unc-3*. Again, we observed region-specific effects. All three markers were ectopically expressed in 10.5 ± 3.7 (mean ± STDV) of the 39 *unc-3*-depleted MNs located in the mid-body region of the VNC (**Fig. 2C-D**). These results are in agreement with a previous study reporting ectopic *ida-1* expression in *unc-3*-depleted MNs [25]. If these ∼11 MNs fully adopt the VC terminal identity, then they should also express genes necessary for acetylcholine biosynthesis since VC neurons are cholinergic. However, this is not the case as expression of *unc-17*/VAChT and *cho-1*/ChT is dramatically affected in *unc-3*-depleted MNs [23]. These data suggest that ∼11 of the 39 *unc-3*-depleted MNs in the mid-body VNC region adopt some, but not all, VC terminal identity features. We will therefore refer to these *unc-3*-depleted MNs as “VC-like” (**Fig. 2G**).

Are the VD-like and VC-like neurons in *unc-3* hermaphrodites distinct populations, or do they represent one population with a mixed (VD-like and VC-like) identity? To test this possibility, we generated *unc-3* hermaphrodites that carry a green fluorescent reporter for VC terminal identity (*ida-1*) and a red reporter for VD identity (*ser-2*). We found no overlap of the two reporers, indicating that the VD-like and VC-like neurons represent two distinct populations (**Fig. 2 E-F**). We further corroborated this result by taking advantage of the invariant lineage and cell body position of all MNs along the *C. elegans* nerve cord (**Suppl. Fig. 2B**). Of note, the VC-like population appears to be lineally related to VC neurons, whereas the VD-like population is not lineally related to VD neurons (**Suppl. Fig. 2B-C**). Lastly, terminal identity markers normally expressed in both VD and VC neurons (*flp-11, ilys-4, twk-46*) display an additive effect in *unc-3* mutants, as they are ectopically expressed in both VD-like and VC-like populations, further suggesting that distinct *unc-3* MN populations express either VD- or VC-specific genes (**Suppl. Fig. 2D-E**).

To summarize, there are 39 *unc-3*-expressing MNs along the mid-body region of the wild-type nerve cord. While loss of *unc-3* uniformly leads to loss of cholinergic identity in all these MNs [23], one population (∼12 MNs) acquires VD-like molecular features, while a second population (∼11 MNs) acquires VC-like molecular features, uncovering a dual role of UNC-3 in these populations (**Fig. 2G**). Of note, the remaining 16 MNs in the VNC of *unc-3* mutants [39 - (12 VD-like + 11 VC-like) = 16] do not gain either VD or VC terminal identity features.

### The dual role of UNC-3 in cholinergic MNs extends to both *C. elegans* sexes

To test whether the dual function of UNC-3 applies to both sexes, we extended our analysis to *C. elegans* males. First, we showed that loss of *unc-3* in males results in loss of several cholinergic MN terminal identity features (**Suppl. Fig. 3A**). Second, we observed ectopic expression of VD-specific terminal identity markers (*oig-1, ser-2*) in 11.9 ± 3.9 (mean ± STDV) out of the 39 *unc-3*-depleted MNs, indicating the presence of “VD-like” neurons in the male nerve cord (**Suppl. Fig. 3B**). Lastly, we asked whether *unc-3* loss leads to ectopic expression of terminal identity markers (*ida-1, srb-16, glr-5*) for male-specific CA neurons. Indeed, we found this to be the case (Suppl. Fig 3C), suggesting the adoption of “CA-like” features by a population of *unc-3*-depleted MNs. Similar to hermaphrodites, these VD-like and CA-like cells were observed in the mid-body region of the male nerve cord.

Taken together, our findings uncover a dual role for UNC-3 in sex-shared cholinergic MNs. UNC-3 is not only required to activate cholinergic MN identity genes [23], but also to prevent expression of terminal identity features of three other nerve cord neuron types (VD, VC, CA). In both sexes, UNC-3 prevents expression of select terminal features of VD neurons in a specific population of cholinergic MNs. In a second population, UNC-3 prevents expression of terminal features normally reserved for sex-specific MNs, i.e., VC terminal features in hermaphrodites and CA terminal features in males. In the ensuing sections, we focus our analysis on *C. elegans* hermaphrodites to dissect the molecular mechanism underlying the dual role of UNC-3.

### UNC-3 is continuously required to prevent expression of VD and VC terminal identity features

Ectopic expression of VC and VD features is observed both at larval and adult stages in *unc-3* null animals (**Fig. 2, Suppl. Fig. 2D-E**). This poses an important question: Is UNC-3 only transiently required during development to, for example, establish a repressive chromatin environment, or is it continuously required to prevent expression of VD and VC terminal identity genes? To address this question, we employed the auxin-inducible degron (AID) system that enables depletion of AID-tagged proteins in a temporally controlled manner [33]. This system requires the tagging of UNC-3 protein with the AID degron fused to a fluorescent reporter gene (mNeonGreen, mNG). When UNC-3::mNG::AID and the plant-specific F-box protein TIR1 are co-expressed in MNs (by crossing animals carrying the *unc-3::mNG::AID* allele with *eft-3::TIR1* transgenic animals), application of the plant hormone auxin on these double transgenic animals induces degradation of UNC-3::mNG::AID (**Fig. 3A-C**). Auxin administration at the L4 stage (last larval stage before adulthood) on *unc-3::mNG::AID*; *eft-3::TIR1* animals resulted in a dramatic depletion of UNC-3 at day 1 adult animals (24 hours after auxin). UNC-3 depletion was accompanied by ectopic expression of VD and VC terminal identity features in nerve cord MNs, demonstrating a post-embryonic requirement for UNC-3 (**Fig. 3D-E**). These findings suggest that UNC-3 is continuously required to prevent expression of VD and VC terminal identity features.

**Figure 3:**
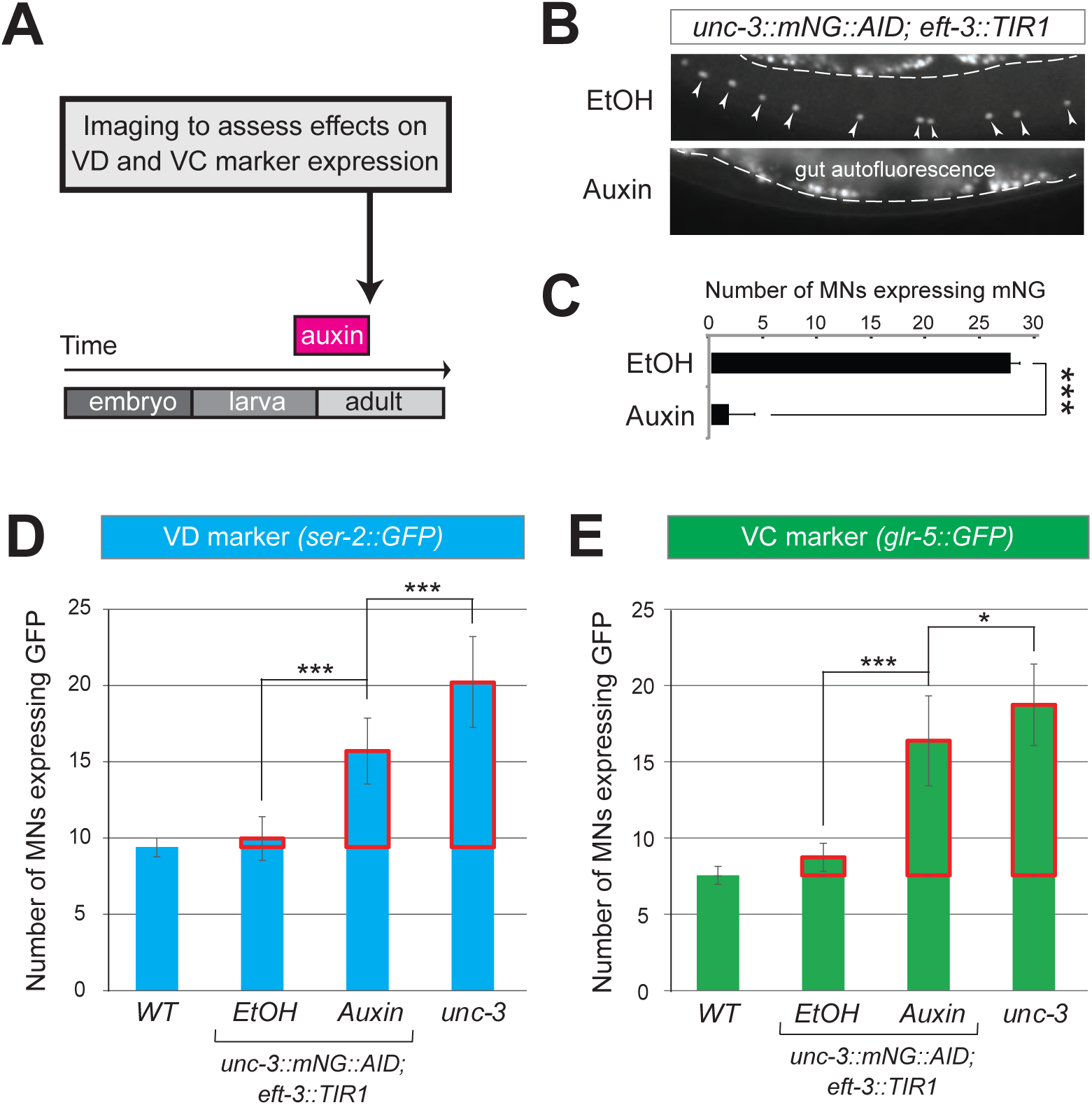
UNC-3 is continuously required to prevent expression of VD and VC terminal identity features. **A:** Schematic showing time window of auxin administration. **B:** Animals of the *unc-3::mNG::AID; eft-3::TIR1* genotype were either administered ethanol (EtOH) or auxin at the L4 stage. Twenty four hours later, expression of endogenous *unc-3* reporter (*unc-3::mNG::AID*) is severely reduced in the nuclei of VNC MNs (arrowheads) at the young adult stage (day 1). The same exact region was imaged in EtOH- and auxin-treated worms. mNG green fluorescent signal is shown in white for better contrast. White dotted line indicates the boundary of intestinal tissue (gut), which tends to be autofluorescent in the green channel. **C:** Quantification of number of MNs expressing the *unc-3::mNG::AID* reporter after EtOH (control) and auxin treatment. N > 12. *** p < 0.001. **D**: Auxin or ethanol (control) were administered at larval stage 3 (L3) on *unc-3::mNG::AID; eft-3::*TIR1 animals carrying the VD marker *ser-2::gfp.* Images were taken at the young adult stage (day 1.5). A significant increase in the number of MNs expressing the VD marker was evident in the auxin-treated animals compared to EtOH-treated controls. For comparison, quantification is provided of *ser-2::gfp* expressing MNs of wild-type animals and *unc-3(n3435)* mutants. Ectopic expression in *unc-3*-depleted MNs is highlighted with a red rectangle. N >20. *** p < 0.001. **E**: Auxin or ethanol (control) were administered at larval stage 4 (L4) on *unc-3::mNG::AID; eft-3::*TIR1 animals carrying the VC marker *glr-5::gfp.* Images were taken at the young adult stage (day 2). A significant increase in the number of MNs expressing the VC marker was evident in the auxin-treated animals compared to EtOH-treated controls. Ectopic expression is highlighted with a red rectangle. N > 11. * p < 0.05; *** p < 0.001.

### UNC-3 acts indirectly to prevent expression of VD and VC terminal identity genes

How does UNC-3 activate cholinergic MN identity genes and simultaneously prevent terminal features of alternative MN identities (e.g., VD, VC) (**Fig. 2G**)? Based on previous reports, the same TF, within the same neuron, can act as a direct activator for a set of genes and a direct repressor for another set of genes [16, 34, 35]. While it is known that UNC-3 acts directly – through its cognate binding site (COE motif) – to activate expression of a large battery of cholinergic MN identity genes, we did not find any COE motifs in the *cis*-regulatory region of VD and VC terminal identity genes (**Table S1**), suggesting an indirect mode of repression via an intermediary factor (**Fig. 4A**). To test this possibility, we focused on VD neurons because, unlike VC neurons, a known activator of VD terminal features has been reported [36–38]. In wild-type animals, the TF UNC-30, ortholog of human PITX1-3, is required to induce VD terminal identity genes. Since UNC-30 is not expressed in cholinergic MNs [38], we hypothesized that UNC-3 prevents expression of UNC-30/PITX (shown as factor X in **Fig. 4A**), leading to inactivation of VD terminal identity genes. However, this is not the case because: (1) ectopic *unc-30* expression is not observed in *unc-3*-depleted MNs, and (2) the ectopic expression of the VD marker (*ser-2*) in *unc-3* mutants was not abolished in *unc-3; unc-30* double mutants (**Suppl. Fig. 4A**). These observations suggest that UNC-3 may act indirectly to prevent expression of VD and VC terminal identity genes through as yet unknown intermediary factors (depicted as X in **Fig. 4A**).

**Figure 4:**
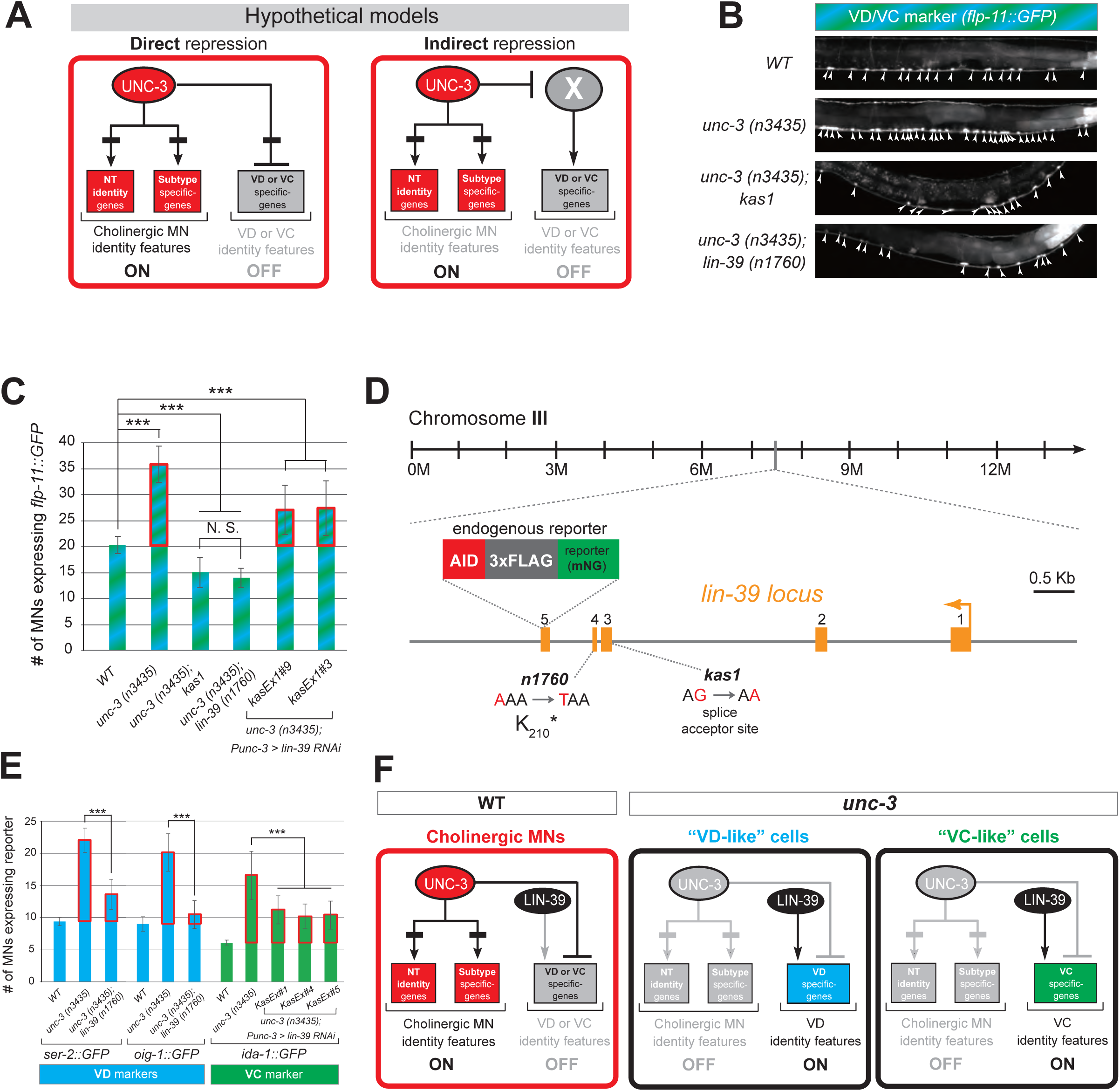
A genetic screen identifies the mid-body Hox protein LIN-39 (Scr/Dfd/Hox4-5) as necessary for ectopic expression of VD and VC terminal features. **A:** Two hypothetical models (direct and indirect) for repression of VD and VC terminal identity genes. **B:** Representative images of L4-stage WT, *unc-3(n3435), unc-3(n3435); kas1*, and *unc-3(n3435); lin-39(n1760)* animals carrying *flp-11::gfp* (VD/VC marker). Arrowheads point to MN cell bodies with *gfp* marker expression. **C**: Quantification graph summarizing results from panel C. The two right-most bars show quantification of two independent transgenic lines driving *lin-39 RNAi* specifically in cholinergic MNs (*Punc-3 > lin-39 RNAi*) of *unc-3 (n3435)* mutants. Ectopic expression in *unc-3*-depleted MNs is highlighted with a red rectangle. N >15. *** p < 0.001. N.S: not significant. **D:** Genetic locus of *lin-39*. Molecular lesions for *kas1* and *n1760* alleles are shown, as well as the *AID::3xFLAG::mNG* cassette inserted at the C-terminus (endogenous reporter). **E**: Quantification of two VD (*ser-2::gfp, oig-1::gfp*) and one VC (*ida-1::gfp*) markers in WT, *unc-3 (n3435), unc-3(n3435); lin-39(n1760)* animals at L4. The three right-most bars show quantification of three independent transgenic lines driving *lin-39 RNAi* specifically in cholinergic MNs (*Punc-3 > lin-39 RNAi*) of *unc-3 (n3435)* mutants. Ectopic expression in *unc-3*-depleted MNs is highlighted with a red rectangle. N > 15. *** p < 0.001. **F**: Schematic that summarizes our findings. Compare panel G to B. *lin-39* is normally expressed in cholinergic MNs but unable to induce expression of VD or VC genes. In the *unc-3* mutant, *lin-39* is now able to induce expression of alternative identity features (VD or VC).

### The mid-body Hox protein LIN-39 (Scr/Dfd/Hox4-5) is the intermediary factor necessary for ectopic expression of VD and VC features in *unc-3* mutants

If our hypothesis is correct (indirect repression model in **Fig. 4A**), mutation of the intermediary factor(s) in the *unc-3* mutant background would selectively eliminate ectopic expression of VD and/or VC terminal identity genes in *unc-3*-depleted MNs. To identify such factor(s), we embarked on an unbiased genetic screen. For the screen, we chose a transgenic *gfp* reporter strain for *flp-11*, an FMRF-like neuropeptide-encoding gene expressed in both VD and VC neurons (**Fig. 1B, Suppl. Fig. 1A**), which is markedly affected by UNC-3 (**Fig. 4B-C, Suppl. Fig. 2D-E**). We mutagenized *unc-3 (n3435); flp-11::gfp* animals with ethyl methanesulfonate (EMS) and visually screened ∼4,200 haploid genomes for mutants in which ectopic *flp-11::gfp* expression in *unc-3*-depleted MNs is suppressed. We isolated one mutant allele (*kas1*) (**Fig. 4B-C**). The phenotype was 100% penetrant as all *unc-3 (n3435); flp-11::gfp* animals carrying *kas1* in homozygosity consistently displayed a dramatic reduction in ectopic *flp-11* expression.

Gross morphological examination of *unc-3 (n3435); kas1*; *flp-11::gfp* hermaphrodites revealed that, unlike *unc-3 (n3435); flp-11::gfp* animals, the introduction of *kas1* is accompanied by a lack of the vulva organ (vulvaless phenotype). Upon a literature survey for TF mutants that are vulvaless, we stumbled across the mid-body Hox gene *lin-39* (ortholog of Dfd/Scr in flies and Hox4-5 in vertebrates) [39, 40], and hypothesized that the molecular lesion of *kas1* may lie in the *lin-39* locus. Indeed, Sanger sequencing uncovered a point mutation on the splice acceptor site (WT: A<u>G</u> > *kas1*: A<u>A</u>) in the second intron of *lin-39* (**Fig. 4D**). Similar to *unc-3 (n3435); kas1* animals, we found that *unc-3 (n3435)* mutants carrying a previously published strong loss-of-function (premature STOP) allele of *lin-39 (n1760)* [40] displayed the same loss of ectopic *flp-11* expression (**Fig. 4B-C**), suggesting that *kas1* is a loss-of-function mutation of *lin-39*. The ectopic expression of *flp-11* in *unc-3(n3435); kas1* animals can be, at least partially, rescued by (1) selective expression of *lin-39* cDNA in cholinergic MNs, and (2) introduction of the *lin-39* wild-type locus in the context of a ∼30kb genomic clone (fosmid) (**Suppl. Fig. 4B**), further corroborating that the *kas1* molecular lesion in the *lin-39* locus is the phenotype-causing mutation.

Because *flp-11* is expressed in both VD and VC neurons, we next tested whether *lin-39* is required for ectopic expression of VD-specific (*ser-2, oig-1*) and VC-specific (*ida-1*) terminal identity genes in *unc-3*-depleted MNs. We found this to be the case by either generating *unc-3 (n3435); lin-39 (n1760)* double mutants (for VD markers) or by performing cholinergic MN-specific RNAi for *lin-39* in *unc-3 (n3435)* animals (for VC marker) (**Fig. 4E**). RNAi was necessary because VC neurons do not survive in *lin-39 (n1760)* animals [41], and the use of the *lin-39* (*n1760)* allele could confound our VC marker quantifications. Of note, all other nerve cord MN subtypes are normally generated in *unc-3 (n3435); lin-39 (n1760)* double mutants [42], indicating that suppression of the *unc-3* phenotype, i.e., loss of ectopic VD or VC gene expression in *unc-3* depleted MNs, is not due to MN elimination in the double mutants. Taken together, our genetic screen identified the mid-body Hox gene *lin-39* to be necessary for ectopic expression of both VD and VC terminal features in *unc-3*-depleted MNs (**Fig. 4F**).

Interestingly, this finding contradicts our initial hypothesis of UNC-3 repressing an intermediary TF in order to prevent expression of VD and VC features (see indirect repression model in **Fig. 4A**) for several reasons. First, *lin-39* is co-expressed with *unc-3* in wild-type cholinergic MNs at the mid-body region of the VNC [22], but is unable to induce VD or VC terminal identity genes (**Fig. 4F**). Second, *lin-39* expression in these MNs is not affected by *unc-3* [22]. Lastly, we found that *lin-39* acts cell-autonomously as cholinergic MN-specific RNAi against *lin-39* in *unc-3 (n3435); flp-11::gfp* animals resulted in a significant reduction of ectopic *flp-11* expression (**Fig. 4C**). In the following Results sections, we provide evidence for an unconventional mechanism through which UNC-3 and LIN-39/Hox select and maintain throughout life key terminal features of cholinergic MNs (**Fig. 4F**).

### UNC-3 prevents a switch in the transcriptional targets of LIN-39 in cholinergic motor neurons

What is the function of LIN-39 in wild-type cholinergic MNs of the VNC? Our previous findings suggested that LIN-39 and UNC-3, together with another mid-body Hox protein, MAB-5 (Antp/Hox6-8) [43], act synergistically to control expression of two terminal identity genes (*unc-129,* ortholog of human BMP; *del-1* / Degenerin-like sodium channel [ortholog of human SCNN1G]) [22]. To test the extent of this synergy, we examined in *lin-39* null animals the expression of 4 additional cholinergic MN terminal identity genes known to be controlled by UNC-3 (*acr-2 / nicotinic acetylcholine receptor; dbl-1 / DPP/BMP-like; nca-1 / sodium channel* [*ortholog of human NALCN*]*, slo-2 / potassium sodium-activated channel* [*ortholog of human KCNT1*]) [23]. In all 4 cases, we found a statistically significant decrease in *lin-39* mutants, and this effect was exacerbated in *lin-39; mab-5* double mutants (**Fig. 5A-B**), indicating that the synergy of LIN-39 with MAB-5 (and UNC-3) extends to multiple terminal identity genes in cholinergic MNs (WT panel in **Fig. 5F**). Of note, while MAB-5 is required for co-activation of cholinergic MN identity genes (**Fig. 5B**), it does not affect the ectopic expression of VD or VC genes observed in *unc-3* mutants (**Fig. 5D-E**).

**Figure 5:**
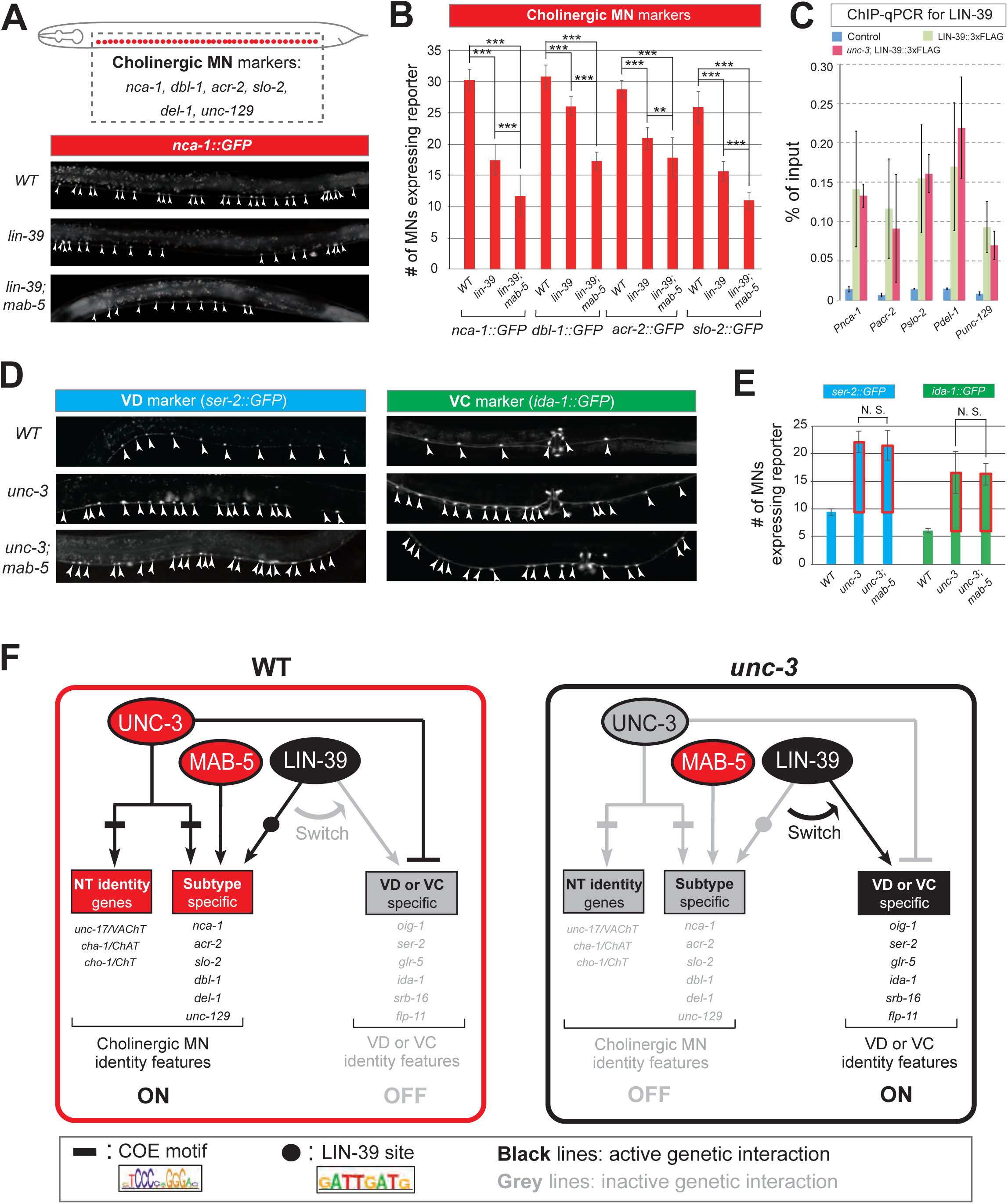
UNC-3 prevents a switch in the transcriptional targets of LIN-39 in cholinergic motor neurons. **A**: Schematic showing cell body position of 39 VNC cholinergic MNs. Markers that label subsets of these MNs are shown *(nca-1, dbl-1, acr-2, slo-2, del-1, unc-129*). Below, representative images are shown of *nca-1::gfp* in *WT, lin-39 (n1760),* and *lin-39 (n1760); mab-5 (1239)* animals at L4 stage. Arrowheads point to MN cell bodies with *gfp* marker expression. Green fluorescence signal is shown in white for better contrast. Dotted black box indicates imaged area. **B**: Quantification of cholinergic MN terminal identity markers (*nca-1, dbl-1, acr-2, slo-2*) *nca-1::gfp* in *WT, lin-39 (n1760),* and *lin-39 (n1760); mab-5 (1239)* animals at L4. N > 15. ** p < 0.01; *** p < 0.001. **C:** ChIP for FLAG (endogenous *lin-39* tagged with *AID::3xFLAG::mNG*) at L3 stage followed by qPCR reveals significant enrichment of LIN-39 binding on its cognate sites at the promoter of multiple cholinergic MN genes (*nca-1, acr-2, slo-2, del-1, unc-129*) when compared to wild-type sample (N2 strain). The endogenous *lin-39* reporter allele (*lin-39::mNG::3xFLAG::AID*) was used for ChIP. **D:** The ectopic expression of VD (*ser-2*) or VC (*ida-1*) terminal identity markers in *unc-3* mutants is not affected by loss of *mab-5*. Strong loss-of-function (null) alleles for *unc-3 (n3435)* and *mab-5 (e1239)* were used. Representative images of L4 stage animals are shown. Arrowheads point to MN cell bodies with *gfp* marker expression. Green fluorescence signal is shown in white for better contrast. **E:** Quantification of data shown in panel D. Ectopic expression is highlighted with a red rectangle. N > 15. N. S: not significant. **F**: Schematic showing the transcriptional switch in LIN-39 targets. In WT animals, UNC-3, MAB-5 and LIN-39 co-activate subtype-specific genes in cholinergic MNs (e.g., *nca-1, dbl-1, unc-129, acr-2*). In *unc-3* mutants, LIN-39 is no longer able to activate these genes, and instead switches to VD- or VC-specific terminal identity genes. Black font: gene expressed. Grey font: gene not expressed. Grey arrows indicate inactive genetic interactions. COE motif (taken from (Kratsios, 2012 #26)) and LIN-39 site (taken from (Weirauch, 2014 #442)) are represented with black rectangles and dots, respectively.

Since UNC-3 controls directly, via its cognate binding site, cholinergic MN terminal identity genes [23], we then asked whether this is the case for LIN-39. Supporting this possibility, we identified via the CIS-BP Database [44] multiple LIN-39 binding sites (previously defined as GATTGATG (Boyle, 2014 #508)) in the *cis*-regulatory region of all 6 aforementioned terminal identity genes (*unc-129, del-1, acr-2, dbl-1, nca-1, slo-2*) (**Table S2**). To assess DNA binding, we performed chromatin immunoprecipitation for endogenous LIN-39 followed by quantitative PCR (ChIP-qPCR) and found evidence for direct binding on at least one site of each terminal identity gene (**Fig. 5C**).

This analysis suggests that LIN-39, similar to UNC-3, regulates directly the expression of multiple terminal identity genes in cholinergic MNs (**Fig. 5F**). Taken together, our data show that, in the absence of UNC-3, the function of LIN-39 in cholinergic MNs is modified. Instead of activating cholinergic MN identity genes, LIN-39 activates VD or VC terminal identity genes in *unc-3*-depleted MNs. Hence, UNC-3 antagonizes the ability of LIN-39 to activate alternative identity genes, thereby preventing a switch in the transcriptional targets of LIN-39 (model schematized in **Fig. 5F**). Interestingly, we found that LIN-39 still binds to its cognate sites on cholinergic MN gene promoters in *unc-3* mutants (**Fig. 5C**), suggesting that LIN-39 is not a rate-limiting factor. That is, there is sufficient amount of LIN-39 in *unc-3*-depleted MNs not only to occupy the LIN-39 sites on cholinergic MN identity genes, but also to induce expression of alternative identity genes.

### LIN-39 is continuously required to control expression of terminal identity genes in cholinergic MNs

The neuronal function of Hox proteins at post-developmental stages is largely unknown [45]. The continuous expression of mid-body Hox *lin-39* in both developing and adult cholinergic MNs led us to investigate whether *lin-39* is required to maintain expression of terminal identity genes in these neurons. To test this idea, we employed clustered regularly interspaced short palindromic repeats (CRISPR)/Cas9-based genome engineering and generated an auxin-inducible *lin-39* allele (*lin-39::mNG::3xFLAG::AID*) that also serves as an endogenous *lin-39* reporter (*mNG*). Animals carrying *lin-39::mNG::3xFLAG::*AID show nuclear mNG expression in MNs located at the mid-body region of the VNC during development and adult stages (**Fig. 6A**), corroborating previous observations with a LIN-39 antibody [46]. Post-embryonic auxin administration on animals carrying the *lin-39::mNG::3xFLAG::AID* resulted in efficient LIN-39 protein depletion that led to a statistically significant reduction in expression of two cholinergic MN identity genes (*acr-2*, *nca-1)* (**Fig. 6A-C, Suppl. Fig. 5A**). We therefore conclude that LIN-39 is continuously required to maintain terminal identity features in cholinergic MNs.

**Figure 6:**
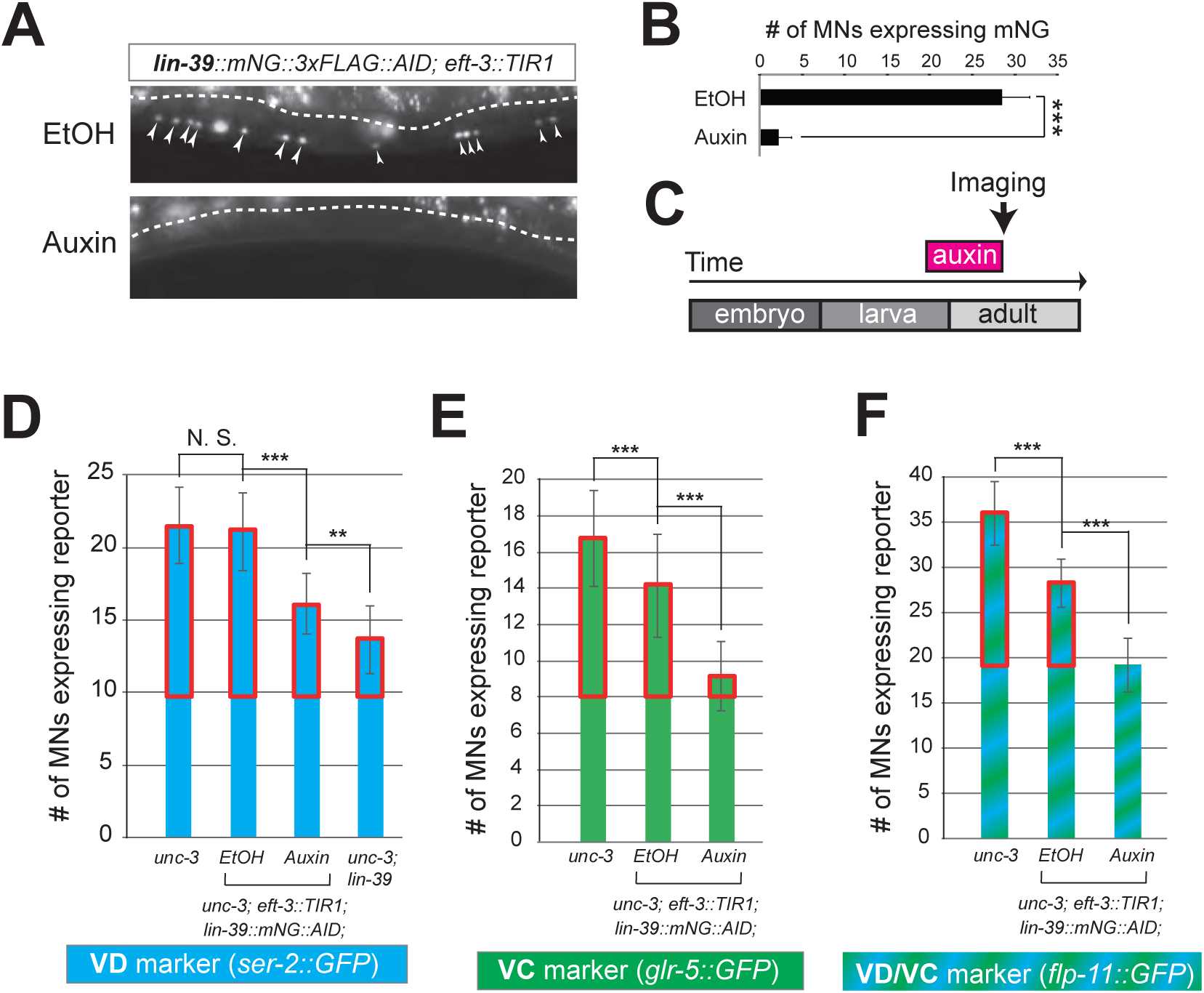
LIN-39 is continuously required to control expression of terminal identity genes. **A:** Animals of the *lin-39::mNG::3xFLAG::AID; eft-3::TIR1* genotype were either administered ethanol (EtOH) or auxin at the L3 stage. Twenty four (24) hours later, expression of endogenous *lin-39* reporter (*lin-39::mNG::3xFLAG::AID*) is severely reduced in the nuclei of VNC MNs (arrowheads) at the young adult stage (day 1). mNG green fluorescent signal is shown in white for better contrast. White dotted line indicates the boundary of intestinal tissue (gut), which tends to be autofluorescent in the green channel. **B:** Quantification of number of MNs expressing the *lin-39::mNG::3xFLAG::AID* reporter after EtOH (control) and auxin treatment. N > 14. *** p < 0.001. **C:** Schematic showing time window of auxin administration. **D-F**: Auxin or ethanol (control) were administered at larval stage 4 (L4) on *unc-3 (n3435); lin-39::mNG::3xFLAG::AID; eft-3::TIR1* animals carrying either the VD marker *ser-2::gfp,* the VC marker *glr-5::gfp,* or the VD/VC marker *flp-11::gfp.* Images were taken at the young adult stage (day 1.6 for *ser-2*, day 1.8 for *glr-5* and day 2 for *flp-11*). A significant increase in the number of MNs expressing the VD marker was evident in the auxin-treated animals compared to EtOH-treated controls. For comparison, quantification of marker expression is also provided in *unc-3 (n3435)* mutants. Ectopic VD and VC expression in *unc-3*-depleted MNs is highlighted with a red rectangle. N > 15. ** p < 0.01, *** p < 0.001, N. S: not significant.

Next, we sought to determine whether LIN-39 is continuously required for the ectopic activation of VD and VC terminal features observed in *unc-3* null animals. Indeed, auxin administration at L4 stage on *unc-3(n3435); lin-39::mNG::3xFLAG::AID* animals carrying either a VD (*ser-2*), VC (*glr-5*), or VD/VC (*flp-11*) marker resulted in a statistically significant suppression of the *unc-3* phenotype when compared to control (treated with ethanol) (**Fig. 6D-F**). We note that hypomorphic effects in the ethanol treated group have been previously reported for other AID-tagged TFs in *C. elegans* [47]. Such effects appear to be target gene-specific, as they were observed for *glr-5* and *flp-11*, but not *ser-2* (**Fig. 6E-F**). Nevertheless, in all three cases we found a statistically significant difference upon auxin administration (compare auxin with EtOH in **Fig. 6D-F**).

To sum up, our findings with the auxin-inducible (**Suppl. Fig. 5A**) and null *lin-39* alleles (**Fig. 4F, 5A-B**) indicate that, in the presence of UNC-3, LIN-39 is required to induce and maintain expression of cholinergic MN terminal identity genes (**Fig. 5F**). In the absence of UNC-3 (**Fig. 6**), LIN-39 is also continuously required - from development and possibly throughout life - for ectopic activation of VD and VC terminal identity genes (**Fig. 5F**).

### LIN-39 is an activator of VD and VC terminal identity genes

The observation that *lin-39* is required for ectopic activation of both VD and VC terminal identity genes in *unc-3*-depleted MNs prompted us to examine the role of *lin-39* in VD and VC neurons of wild-type animals. Does LIN-39 control the same VD- and VC-specific terminal identity genes that become ectopically expressed in *unc-3* mutants?

To this end, we leveraged our endogenous *lin-39* reporter (*lin-39::mNG::3xFLAG::AID*) to assess expression in wild-type VD neurons at the mid-body region of the VNC, and found this to be the case (**Suppl. Fig. 5C**). Next, we found that LIN-39 is required to induce expression of VD terminal identity genes (*ser-2, oig-1*) (**Fig. 7A-B**). To gain further mechanistic insights, we then asked whether *lin-39* acts together with UNC-30, the known activator of GABAergic MN identity genes (introduced earlier). Apart from confirming previous observations of UNC-30 controlling the VD-specific *oig-1* gene [36, 48], we also found that *ser-2* (**Fig. 7B**) and *flp-11* (**Suppl. Fig. 5C**) constitute novel UNC-30 targets in VD neurons. To test for synergistic effects, we generated *lin-39; unc-30* double mutants and observed stronger effects on *ser-2* and *flp-11* expression than either single mutant (**Fig. 7B, Suppl. Fig. 5C**). Such additive effects indicate that *lin-39* and *unc-30* act in parallel to co-activate VD terminal identity genes. Importantly, expression of other UNC-30 targets in GABAergic MNs, such as *flp-13* (DD-specific terminal identity marker) [36, 49, 50] and genes expressed in both DD and VD neurons (*unc-25/GAD*, *unc-47/VGAT*), is unaffected in *lin-39* mutants (**Fig. 7B, Suppl. Fig. 5E**). Unlike UNC-30 that broadly controls multiple terminal features in VD neurons, we conclude that *lin-39* is selectively required for activation of VD-specific terminal identity genes (**Fig. 7I**). To test for a maintenance role in VD neurons, we administered auxin at a post-developmental stage (L3) on animals carrying the *lin-39::mNG::3xFLAG::AID* allele. We found that LIN-39 is continuously required to maintain expression of the VD terminal identity gene *ser-2* (**Fig. 7C**).

**Figure 7:**
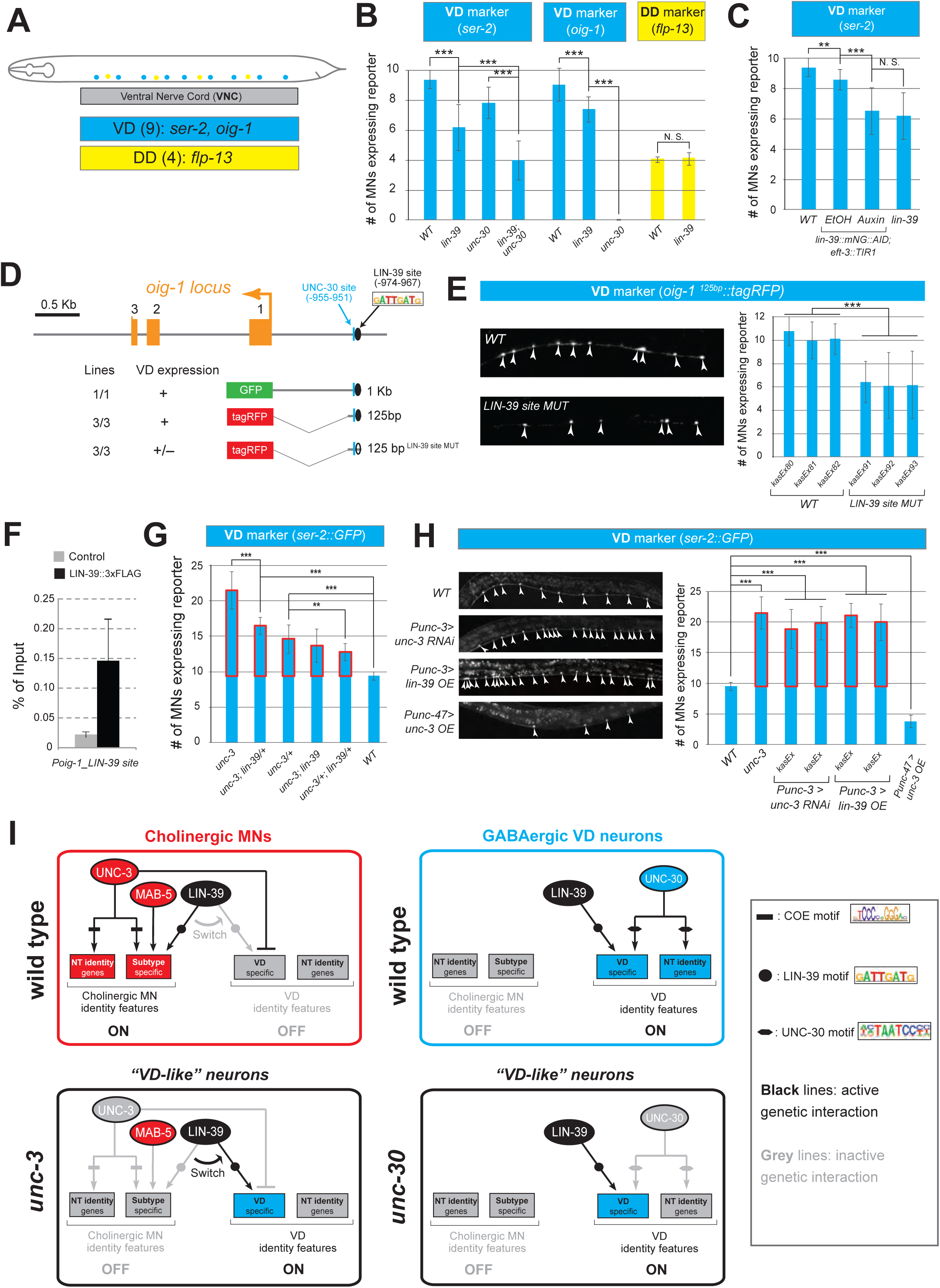
LIN-39 is an activator of VD terminal identity genes. **A:** Schematic showing 9 VD and 4 DD neurons populating the VNC. In addition, 4 VD and 2 DD neurons are located in ganglia flanking the VNC (not shown because they were excluded from our analysis). **B:** Quantification of two VD (*ser-2::gfp, oig-1::gfp*) and one DD (*flp-13::gfp*) markers in WT and *lin-39 (n1760)* animals at L4. Both VD markers were also tested in *unc-30 (e191)* mutants. Double *lin-39 (n1760); unc-30 (e191)* mutants showed a more severe reduction in expression of the VD marker *ser-2::gfp* compared to each single mutant. N > 15. *** p < 0.001. N. S: not significant. **C:** Auxin or ethanol (control) were administered at larval stage 3 (L3) on *lin-39::mNG::3xFLAG::AID; eft-3::TIR1* animals carrying the VD marker *ser-2::gfp.* Images were taken at the young adult stage (day 1.5). A significant decrease in the number of MNs expressing the VD marker was evident in the auxin-treated animals compared to EtOH-treated controls. For comparison, quantification of marker expression is also provided in WT and *lin-39 (n1760)* animals. N > 15. ** p < 0.01, *** p < 0.001. N. S: not significant. **D**: Schematic of *oig-1* locus showing location of LIN-39 and UNC-30 sites. Transgenic reporter lines carrying either 1kb or 125bp elements of the *oig-1* promoter show expression in VD neurons. **E**: Mutation of a single *lin-39* site in the context of transgenic animals carrying the 125bp *oig-1* element (*oig-1_125bp ^lin-39^ ^site^ ^MUT^::tagRFP*) results in significant reduction in the number of VD neurons expressing the reporter at L4. Representative images are shown on the left with arrowheads pointing to MN cell bodies with marker expression. Red fluorescence signal is shown in white for better contrast. Quantification of three independent transgenic lines is provided on the right. N = 14. *** p < 0.001. **F**: ChIP for FLAG (endogenous *lin-39* tagged with *AID::3xFLAG::mNG*) at L3 stage followed by qPCR in wild-type (N2 strain) and *lin-39::mNG::3xFLAG::AID* animals. Compared to control (N2 strain), significant enrichment of LIN-39 binding on the LIN-39 site of the *oig-1* promoter was observed. **G:** Quantification of the VD marker (*ser-2::gfp*) in *unc-3 (n3435), unc-3 (n3435); lin-39 (n1760)/+, unc-3 (n3435)/+, unc-3 (n3435); lin-39 (n1760), unc-3 (n3435)/+; lin-39 (n1760)/+,* and WT animals at L4. Ectopic expression is highlighted with a red rectangle. N > 15. ** p < 0.01, *** p < 0.001. **H:** Representative images of the VD marker (*ser-2::gfp*) expression on the left in L4 stage transgenic animals that either down-regulate *unc-3* in cholinergic MNs (*Punc-3 > unc-3 RNAi*), over-express *lin-39* in cholinergic MNs (*Punc-3 > lin-39 OE*), or over-express *unc-3* in VD neurons (*Punc-47 > unc-3 OE*). Arrowheads point to MN cell bodies with *gfp* marker expression. Green fluorescence signal is shown in white for better contrast. Quantification is provided on the right. Ectopic expression is highlighted with a red rectangle. Two independent transgenic lines were used for *Punc-3 > unc-3 RNAi* and *Punc-3 > lin-39 OE.* N > 13. *** p < 0.001. **I**: Schematic that summarizes the function of LIN-39 in WT cholinergic MNs, *unc-3*-depleted MNs (“VD-like” neurons), and GABAergic VD neurons. COE motif is taken from [23]. LIN-39 site is taken from [44]. UNC-30 site is taken from [50].

Does *lin-39* control expression of terminal identity genes in sex-specific VC neurons? We used the auxin-inducible *lin-39::mNG::3xFLAG::AID* allele to address this question because VC neurons do not survive in *lin-39 (n1760)* null animals [41]. We applied auxin at a late larval stage (L3-L4) to knock-down LIN-39 and observed that VC neurons do not die, providing an opportunity to test for putative effects on VC terminal identity gene expression. Indeed, we found a statistically significant reduction in the number of VC neurons expressing *srb-16* (compare auxin and ethanol in **Suppl. Fig. 5B**).

Taken together, *lin-39* is required for expression of VD- and VC-specific terminal identity genes. In VD neurons, LIN-39 acts synergistically with UNC-30/PITX to co-activate expression of VD-specific genes (top right panel in **Fig. 7I**). Collectively, these findings on VD and VC neurons together with observations on cholinergic MNs (**Fig. 5A-B**) show that, in different MN subtypes, the mid-body Hox gene *lin-39* controls expression of distinct terminal identity genes, likely due to synergy with distinct TFs (i.e., UNC-3 and MAB-5 in cholinergic MNs versus UNC-30 in VD neurons [also summarized in **Fig. 7I**]).

### LIN-39 directly activates the VD terminal identity gene *oig-1*

Does LIN-39 act directly or indirectly to activate VD and VC terminal identity genes? To this end, we interrogated the *cis*-regulatory regions of VD, VC, and VD/VC terminal identity genes for the presence of LIN-39 binding sites. We indeed found several sites in all these genes (**Table S2**), indicating direct regulation. To test the functionality of these putative LIN-39 binding sites, we honed in on the VD-specific terminal identity gene *oig-1*. A previous study identified a minimal 125bp *cis*-regulatory element upstream of *oig-1* as sufficient to drive reporter gene expression in VD neurons [48] (**Fig. 7D**). We independently confirmed this observation, and further found that the 125bp element contains a single LIN-39 site. Mutation of this site in the context of transgenic *oig-1* reporter animals (*oig-1 ^125bp^ ^LIN-39^ ^site^ ^MUT^::tagRFP*) leads to a significant reduction of tagRFP expression in VD neurons (**Fig. 7E**), phenocoping the effect observed in *lin-39 (n1760)* null mutants (**Fig. 7B**). Next, we performed ChIP-qPCR using primers flanking the LIN-39 site within the 125bp element and indeed confirmed LIN-39 binding (**Fig. 7F**). We conclude that LIN-39 acts directly, by recognizing its cognate site, to activate expression of the VD-specific gene *oig-1*. Interestingly, a functional binding site for UNC-30/PITX also exists in this 125bp element [48, 50], and is spaced 11 base pairs apart from the LIN-39 site (**Fig. 7D**), indicating that LIN-39 and UNC-30 control *oig-1* by recognizing distinct and in close proximity *cis*-regulatory motifs. Lastly, we surveyed available ChIP-Seq data for LIN-39 from the modENCODE project [51] using WormBase (https://wormbase.org) and found evidence for LIN-39 binding in the *cis*-regulatory region of *ida-1* (VC identity marker) and *ilys-4* (VD/VC identity marker) (**Suppl. Fig. 6**), suggesting that the mode of direct regulation by LIN-39 likely extends beyond *oig-1*.

### The LIN-39-mediated transcriptional switch appears unidirectional

The above analysis on *oig-1* indicates that LIN-39 and UNC-30/PITX directly co-activate expression of VD-specific genes (top right panel in **Fig. 7I**). Similarly, LIN-39 and UNC-3 directly co-activate terminal identity genes in cholinergic MNs (top left panel in **Fig. 7I**). Since the absence of UNC-3 leads to ectopic activation of VD-specific genes, we next considered the converse possibility: Does the absence of UNC-30/PITX lead to ectopic activation of cholinergic MN terminal identity genes in GABAergic VD neurons? However, this appears not to be the case as expression of 4 cholinergic MN markers *(acr-2, slo-1, unc-129, del-1),* normally co-activated by UNC-3 and LIN-39 (**Fig. 5A-B**), is unaffected in *unc-30* mutants (**Suppl. Fig. 7**), suggesting that UNC-30/PITX, unlike UNC-3, does not exert a dual role in GABAergic VD neurons.

These findings indicate that the LIN-39-mediated transcriptional switch is unidirectional (summarized in **Fig. 7I**). In the absence of UNC-3, LIN-39 is no longer able to activate cholinergic MN terminal identity genes and instead switches to VD-specific genes (“VD-like” neurons in **Fig. 7I**). However, LIN-39 is not able to switch targets (from VD- to cholinergic MN-terminal identity genes) in the absence of UNC-30/PITX.

### The LIN-39-mediated transcriptional switch depends on UNC-3 and LIN-39 levels

How does the absence of UNC-3 lead to ectopic and *lin-39*-dependent activation of VD terminal identity genes in cholinergic MNs (**Fig. 7I**)? In principle, UNC-3 and LIN-39 could physically interact in order to co-activate expression of cholinergic MN terminal identity genes. In the absence of *unc-3*, this interaction would be disrupted and LIN-39 becomes available, in cholinergic MNs, to assume its VD function, i.e., to activate VD-specific terminal identity genes (**Fig. 7I**). However, our co-immunoprecipitation (co-IP) experiments on UNC-3 and LIN-39 in a heterologous system (HEK cells) did not provide evidence for physical interaction (**Suppl. Fig. 8**).

Next, we hypothesized that the levels of UNC-3 and LIN-39 may play a critical role in the observed LIN-39-mediated transcriptional switch (**Fig. 7I**). Indeed, we found a gene dosage relationship between *unc-3* and *lin-39*. Loss of one *unc-3* copy (*unc-3 (n3435)/+*) caused slight ectopic expression of VD genes (*ser-2* in **Fig. 7G** and *flp-11* in **Suppl. Fig. 9A**), but that ectopic expression is decreased by loss of one *lin-39* copy (*unc-3 (n3435)/+; lin-39 (n1760)/+*) (**Fig. 7G, Suppl. Fig. 9A**). Accordingly, loss of one *lin-39* copy in *unc-3* null animals *(unc-3 (n3435); lin-39 (n1760)/+)* also reduced, but did not eliminate, ectopic expression of VD genes (**Fig. 7G, Suppl. Fig. 9A**). These data firmly suggest there is a close stoichiometric relationship between UNC-3 and LIN-39. Supporting this scenario, knock-down of *unc-3* with RNAi specifically in cholinergic MNs also led ectopic expression of VD genes (**Fig. 7H, Suppl. Fig. 9B-C**), whereas overexpression of UNC-3 in VD neurons resulted in repression of VD gene expression (**Fig. 7H, Suppl. Fig. 9B-C**). Lastly, we asked whether LIN-39 is sufficient to induce expression of VD terminal identity genes in cholinergic MNs. Indeed, we found this to be the case upon over-expression of LIN-39 using cholinergic MN gene promoters (**Fig. 7H, Suppl. Fig. 9D-E**). In conclusion, we propose that the LIN-39-mediated transcriptional switch observed in *unc-3* mutants critically depends on UNC-3 and LIN-39 levels (left panels in **Fig. 7I**).

### Ectopic expression of VD terminal identity genes in cholinergic motor neurons is associated with locomotion defects

The dual role of UNC-3 revealed by our molecular analysis (**Fig. 2**) led us to posit that the severe locomotion defects observed in *unc-3* animals may represent a composite phenotype [27, 52]. In other words, these defects are not only due to loss of expression of cholinergic MN terminal identity determinants (e.g., *unc-17*/VAChT, *cha-1*/ChAT, *del-1*/Degenerin-like sodium channel, *acr-2*/acetylcholine receptor [ortholog of CHRNE]), but also due to the ectopic expression of VD and VC terminal features (e. g., *ser-2*/serotonin receptor [ortholog of HTR1D], *flp-11*/FRMR-like neuropeptide, *glr-5*/Glutamate receptor [ortholog of GRID], *srb-16*/GPCR) in *unc-3-*depleted MNs. To genetically separate these distinct molecular events, we generated *unc-3 (n3435); lin-39 (n1760)* double mutants, which do display loss of cholinergic MN terminal identity genes, but the ectopic expression of VD and VC terminal features is suppressed (**Fig. 4F**). We predicted that if ectopic expression of VD and VC genes contributes to locomotion defects, then *unc-3 (n3435)* mutants would display more severe locomotion defects than *unc-3 (n3435); lin-39 (n1760)* double mutants. To test this, we performed high-resolution behavioral analysis of freely moving adult (day 1) *C. elegans* animals using automated multi-worm tracking technology [52, 53]. This analysis can quantitate multiple features related to *C. elegans* locomotion (*e.g.,* speed, crawling amplitude, curvature, pause, forward and backward locomotion) and, most importantly, each feature can be localized to a specific part of the nematode’s body (e. g., head, mid-body, tail). Since *unc-3* and *lin-39* expression uniquely overlaps in mid-body nerve cord MNs that innervate mid-body muscles, we hypothesized that loss of *unc-3* and/or *lin-39* genes would have effects on mid-body posture and motion, and thereby focused our analysis on mid-body curvature features. Of the 49 mid-body features examined, 29 were significantly different in *unc-3* single mutants when compared to wild-type (N2 strain) animals (see **Table S3** for all 49 features). Intriguingly, 12 of these 29 features (41.37%) were significantly suppressed in *unc-3; lin-39* double mutants (**Fig. 8A, Suppl. Fig. 10A, Table S3**), suggesting that suppression of these behavioral defects could be attributed to suppression of the ectopically expressed VD and VC terminal identity genes in these double mutants. We found no evidence for suppression of the remaining 17 features in *unc-3; lin-39* double mutants, likely due to the fact that UNC-3 also controls other terminal identity genes, such as NT pathway genes (**Fig. 7I**), independently of LIN-39.

**Figure 8:**
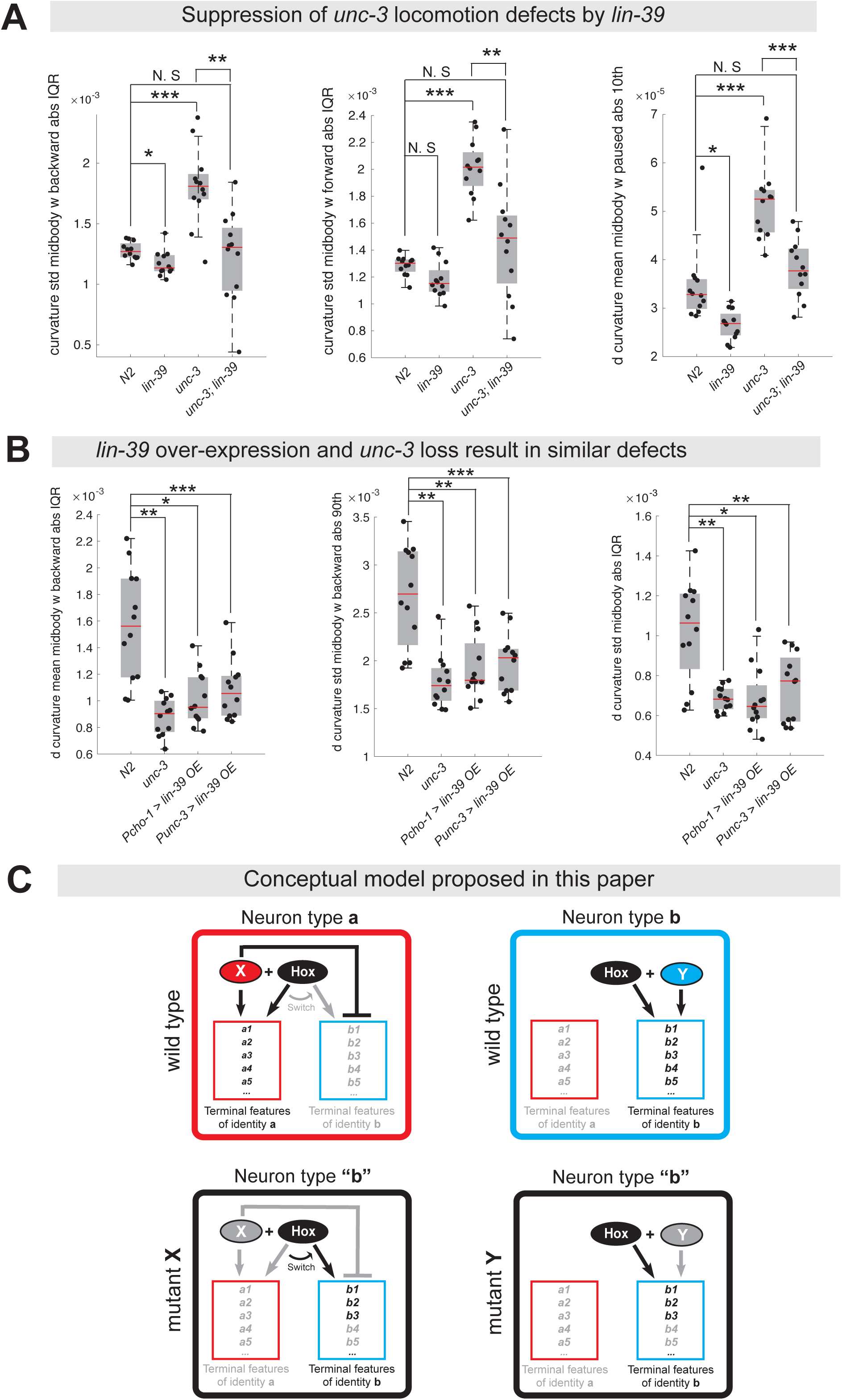
Ectopic expression of VD terminal identity genes in cholinergic motor neurons is associated with locomotion defects. **A**: Examples of three mid-body locomotion features that are significantly affected in *unc-3 (n3435)* animals, but markedly improved in *unc-3 (n3435); lin-39 (n1760)* double mutant animals. Each black dot represents a single adult animal. The unit for the first two graphs is 1/microns. The unit for the graph on the right is 1/(microns*seconds). N = 12. Additional mid-body features affected in *unc-3 (n3435)* animals, but improved in *unc-3 (n3435); lin-39 (n1760)* mutants are provided in **Suppl. Figure 10A** and **Table S3**. * p < 0.01, ** p < 0.001, *** p < 0.0001. **B**: Examples of three mid-body locomotion features affected in *unc-3 (n3435)* mutants and animals over-expressing *lin-39* in cholinergic MNs. Each black dot represents a single adult animal. The unit for the Y axis is 1/(microns*seconds). N = 12. Additional mid-body features affected in *unc-3 (n3435)* and *lin-39* over-expressing animals are provided in **Suppl. Figure 10B-C** and **Table S3**. * p < 0.01, ** p < 0.001, *** p < 0.0001. **C:** Conceptual model summarizing the findings of this paper. Grey font: not expressed gene. Black font: expressed gene. Grey arrow: inactive genetic interaction. Black arrow: active genetic interaction.

Next, we asked whether ectopic expression of VD terminal identity genes in otherwise wild-type animals can lead to locomotion defects. To test this, we took advantage of our transgenic animals that selectively over-express LIN-39 in cholinergic MNs (*Pcho-1 > LIN-39, Punc-3 > LIN-39*) (**Fig. 7H, Suppl. Fig. 9D**). First, we confirmed that in these animals expression of cholinergic MN terminal identity genes is unaffected (**Suppl. Fig. 10D**). Second, we found that LIN-39 overexpression led to ectopic activation of VD, but not VC, terminal identity genes in cholinergic MNs (**Suppl. Fig. 10D**), providing an opportunity to specifically assess the consequences of ectopic VD gene expression on animal locomotion. We found that 9 of the 29 (31.03%) mid-body features affected in *unc-3 (n3435)* animals were also altered in animals over-expressing *lin-39* in cholinergic MNs (**Fig. 8B, Suppl. Fig. 10B-C, Table S3**).

In conclusion, our behavioral analysis is in agreement with our molecular findings. At the molecular level, we found that *lin-39*/Hox is necessary for the ectopic expression of VD terminal identity genes in *unc-3* mutants. At the behavioral level, this *lin-39*-dependent, ectopic expression of terminal identity genes is accompanied by locomotion defects.

## DISCUSSION

During development, individual neuron types must select their unique terminal identity features, such as expression of NT receptors, ion channels and neuropeptides. Continuous expression of these features - from development through adulthood - is essential for safeguarding neuronal terminal identity, and thereby ensuring neuronal function [1, 3, 4]. Here, we provide critical insights into the mechanisms underlying selection and maintenance of neuron type-specific terminal identity features by using the well-defined MN subtypes of the *C. elegans* nerve cord as a model. First, we report that, in cholinergic MNs, the terminal selector-type TF UNC-3 has a dual role; UNC-3 is not only required to promote cholinergic MN identity features, but also to prevent expression of multiple terminal features normally reserved for three other ventral cord neuron types (VD, VC, CA). Second, we provide evidence that cholinergic MNs can secure their terminal identity throughout life by continuously relying on UNC-3’s dual function without the need to employ an entirely new maintenance program. Third, we propose an unconventional mechanism underlying the dual function of a neuron type-specific TF, as we find UNC-3 necessary to prevent a switch in the transcriptional targets of the mid-body Hox protein LIN-39 (Scr/Dfd/Hox4-5) (**Fig. 7I**). Lastly, our findings shed light upon the poorly explored post-embryonic function of Hox proteins in the nervous system by uncovering that LIN-39/Hox is continuously required to maintain expression of multiple terminal identity genes in MNs.

Is there a need for mechanisms that continuously prevent expression of alternative identity features in a post-mitotic neuron? Or, such mechanisms become superfluous once neurons have restricted their developmental potential by committing to a specific terminal identity? This fundamental question remains unanswered, in part due to the fact that most neuron type-specific TFs with a dual function have been studied during embryonic stages. Our auxin experiments uncovered a continuous requirement for the dual role of UNC-3. Post-embryonic depletion of UNC-3 not only results in failure to maintain cholinergic MN terminal features [23], but is also accompanied by ectopic expression of alternative identity features (e.g., VD, VC). These findings reveal a simple and economical mechanism that secures neuronal terminal identity features: the same TF is continuously required - from development throughout life - to not only activate neuron type-specific identity genes, but also prevent expression of “unwanted” terminal features. A continuous requirement for repression of terminal identity genes has been previously reported for three other TFs, *bnc-1*/BNC and *mab-9*/Tbx20 in *C. elegans* MNs and *ewg*/NRF1 in the *Drosophila* eye [47, 54]. However, it is currently unknown whether these factors exert a dual and continuous function like UNC-3.

Numerous cases of neuron type-specific TFs with a dual role have been previously described in both vertebrate and invertebrate models systems [8–15]. Although the underlying mechanisms often remain unclear, recent studies proposed two modes of action. On the one hand, such TFs can act directly to activate “desired” terminal identity features and repress (also directly) “unwanted” features. On the other hand, neuron type-specific TFs can also act indirectly. For example, suppression of alternative identity features can be achieved by repressing an intermediary TF that normally activates such features [11]. Here, we reveal an alternative mechanism by probing the dual function of UNC-3 (**Fig. 7I, 8F**). Activation of cholinergic MN identity features relies on synergy between UNC-3 and two mid-body Hox proteins, LIN-39 and MAB-5. Intriguingly, suppression of alternative terminal identity genes (e.g., VD, VC) is not achieved directly by UNC-3, i.e., via its cognate binding site. This contrasts the previously described function of UNC-3 as direct repressor of terminal identity genes in the chemosensory ASI neurons of *C. elegans* [55]. We further found no evidence of UNC-3 repressing an intermediary TF (**Fig. 4A**). Instead, UNC-3 is necessary to prevent a switch in the transcriptional targets of LIN-39/Hox. Upon genetic removal of *unc-3*, LIN-39 is no longer able to co-activate cholinergic MN terminal identity genes and switches its function to induce expression of alternative identity (e.g., VD, VC) features (**Fig. 7I**). However, the precise molecular mechanism of how UNC-3 prevents this LIN-39-mediated transcriptional switch remains poorly understood. A recent study proposed that UNC-3 is able to restrict cellular plasticity, at least partially by organizing chromatin and regulating the distribution of H3K9 methylation, a transcriptional repression mark [56]. We therefore favor a model in which the presence of UNC-3 may lead to H3K9 methylation of VD and VC gene loci. In the absence of UNC-3, this repression mark is compromised, enabling LIN-39 to assume its wild-type function, i.e., to activate VD- or VC-specific terminal identity genes. This hypothetical model is further supported by studies on Ebf1 (UNC-3 ortholog) in B cells [57].

A large body of work in multiple model systems has established that Hox proteins are required, during early development, for neuronal diversity, cell survival, axonal pathfinding and circuit assembly [29, 30, 58–64]. However, the function and downstream targets of Hox proteins during post-embryonic stages are largely unknown. Our contributions towards this knowledge gap are twofold. First, the auxin experiments revealed that the mid-body Hox protein LIN-39 is continuously required to control expression of MN terminal identity genes. Second, we uncovered multiple terminal identity genes as downstream targets of LIN-39 in different MN subtypes (cholinergic MNs: *acr-2, dbl-1, nca-1, slo-2;* VD neurons: *oig-1, ser-2, flp-11;* VC neurons: *srb-16)*. Given the maintained expression of Hox genes in the adult nervous system of flies, mice and humans [45, 58, 65], our findings may be broadly transferable.

TFs able to broadly activate many distinct terminal identity features of a specific neuron type (e.g., NT biosynthesis components, NT receptors, ion channels, neuropeptides) have been termed “terminal selectors” [2]. Several dozens of terminal selectors have been described thus far in multiple model systems including worms, flies and mice [3, 4, 28]. However, whether terminal selectors are required to prevent expression of alternative identity features is not known. Our findings on the terminal selector UNC-3 suggest that this is the case, and further reveal a continuous requirement for prevention of such features by UNC-3. In the future, it will be interesting to see whether the lessons learnt from UNC-3 broadly apply to other terminal selectors described across species. Supporting this intriguing possibility, Pet-1, the terminal selector of mouse serotonergic neurons has been recently shown to repress the terminal identity gene *Hypocretin receptor 1* (*Hcrtr1*) [34].

Although our study employs an extensive repertoire of terminal identity markers for distinct MN subtypes, the extent of alternative identity features (e.g., VD, VC) being ectopically expressed in *unc-3*-depleted MNs remains unknown. Future unbiased transcriptional profiling of *unc-3-*depleted MNs could help address this issue. In addition, the strong axonal defects in MNs of *unc-3* mutants preclude any further attempts to assess whether the observed VD-like and VC-like cells, as defined by molecular markers, also acquire morphological features of VD and VC neurons, respectively [26]. Nevertheless, the examination of multiple MN terminal identity markers at single-cell resolution enabled us to make an interesting observation: Although all *unc-3*-depleted nerve cord MNs uniformly lose their cholinergic identity, one subpopulation acquires VD terminal features (“VD-like” neurons) and another subpopulation acquires VC terminal features (“VC-like” neurons). This intriguing observation may be analogous to findings described in the mammalian neocortex, where genetic removal of the TF *Satb2* leads to loss of pyramidal neuron identity (UL1 subtype), and concomitant gain of molecular features specific to two other pyramidal neuron subtypes (DL, UL2) [10]. Apart from supporting the notion that neuron type-specific TFs often suppress features of functionally related neuronal subtypes [31], the cases of UNC-3 and Satb2 highlight that genetic removal of a single TF in a seemingly homogeneous neuronal population (UNC-3 in cholinergic MNs, Satb2 in UL1 pyramidal neurons) can lead to nonuniform outcomes, i.e., activation of several alternative identity programs.

One important aspect of our findings relates to the experiments performed on both *C. elegans* sexes. The organization of sex-shared and sex-specific MNs in the *C. elegans* nerve cord is reminiscent of MNs found in other invertebrate and vertebrate nervous systems. However, it is largely unknown whether and how acquisition of terminal identity in sex-shared MNs is coordinated with prevention of sex-specific MN identity features. The availability of terminal identity markers for sex-specific MNs (VC in hermaphrodites, CA in males) in *C. elegans* enabled us to discover that, in sex-shared cholinergic MNs, UNC-3 is continuously required to suppress terminal features normally reserved for sex-specific MNs. Given that the *unc-3* ortholog Ebf2 is co-expressed with Hox genes in sex-shared cholinergic MNs of the mouse spinal cord [22, 66], the molecular mechanism described here for suppression of “unwanted” terminal identity features (e.g., sex-specific MN features) by UNC-3 may be conserved across species.

From an evolutionary standpoint, our findings highlight the employment of economical solutions to evolve novel cell types in the nervous system. The same Hox protein (LIN-39) synergizes with distinct terminal selectors in different MNs. In GABAergic (VD) neurons, LIN-39 synergizes with UNC-30/PITX to control expression of VD terminal identity genes, whereas in cholinergic MNs LIN-39 synergizes with UNC-3 to control cholinergic MN identity genes (**Fig. 7I**). We speculate that the *unc-3* mutant “state” may constitute the “ground state”. For example, the “VD-like” neurons in *unc-3* mutants that express LIN-39 may represent an ancient cell type that was altered to become a new cell type through the recruitment of distinct terminal selectors (**Fig. 7I**, and conceptual model in **Fig. 8C**). Hence, the amount of genetic information required for evolution of new cell types is kept to minimum. The recruitment of UNC-30/PITX enabled “VD-like” cells to fully adopt GABAergic VD neuron terminal identity, as evident by the ability of UNC-30/PITX to control expression of GABA synthesis proteins [37, 38] (**Fig. 7I**). Similarly, recruitment of UNC-3 enabled “VD-like” cells to become cholinergic MNs. In this “new” cholinergic cell type, UNC-3 not only antagonizes the ability of LIN-39 to activate VD-specific genes, but also synergizes with LIN-39 to co-activate cholinergic MN terminal identity genes. We hope the strategy described here of a terminal selector preventing a Hox transcriptional switch may provide a conceptual framework for future studies on terminal identity and evolution of neuronal cell types.

## ACKNOWLEDGEMENTS

We thank the Caenorhabditis Genetics Center (CGC), which is funded by NIH Office of Research Infrastructure Programs (P40 OD010440), for providing strains. We thank Anthony Osuma, Melanie Le Gouez, and Minhkhoi Nguyen for generating *lin-39* and *oig-1* reporter strains. We are grateful to Oliver Hobert, Robert Carillo, Catarina Catela, and Daniele Canzio for comments on this manuscript. This work was funded by an NINDS grant (R00NS084988) and a Whitehall Foundation grant to P.K.

## MATERIALS AND METHODS

### C. elegans strains

Worms were grown at 15°C, 20°C or 25°C on nematode growth media (NGM) plates seeded with bacteria (*E.coli* OP50) as food source [27]. Mutant alleles used in this study: *unc-3 (n3435) X, lin-39 (n1760) III, mab-5 (e1239) III, unc-30 (e191) IV, him-5 (e1490) V, him-8 (e1489) IV, lin-39 (kas1) III.* CRISPR-generated alleles: *unc-3 (ot837 [unc-3::mNG::AID]) X, lin-39 (kas9 [lin-39::mNG::AID]) III*.

RNAi and overexpression transgenes used in this study:

> *kasEx68 [punc-3_558bp>lin-39 RNAi#3 + myo-2::GFP],*
>
> *kasEx69 [punc-3_558bp>lin-39 RNAi#9 + myo-2::GFP],*
>
> *kasEx70 [punc-3_558bp>lin-39 RNAi#1 + myo-2::GFP],*
>
> *kasEx71 [punc-3_558bp>lin-39 RNAi#4 + myo-2::GFP],*
>
> *kasEx72 [punc-3_558bp>lin-39 RNAi#5 + myo-2::GFP],*
>
> *kasEx73 [punc-3_558bp>unc-3 RNAi#6 + myo-2::GFP],*
>
> *kasEx74 [punc-3_558bp>unc-3 RNAi#8 + myo-2::GFP],*
>
> *kasEx35 [punc-3_558bp>lin-39 cDNA OE#1 + myo-2::GFP],*
>
> *kasEx36 [punc-3_558bp>lin-39 cDNA OE#2 + myo-2::GFP],*
>
> *kasEx37 [punc-3_558bp>lin-39 cDNA OE#5 + myo-2::GFP],*
>
> *kasEx75 [punc-47>unc-3 cDNA + myo-2::GFP],*
>
> *kasEx76 [punc-3_558bp>lin-39 cDNA#2.2 + myo-2::GFP],*
>
> *kasEx77 [punc-3_558bp>lin-39 cDNA#10.1 + myo-2::GFP],*
>
> *kasEx33 [lin-39 fosmid WRM0616aE11#32 + myo-2::GFP],*
>
> *kasEx34 [lin-39 fosmid WRM0616aE11#34 + myo-2::GFP],*
>
> *kasEx78 [punc-3_558bp>unc-3 RNAi#n1’ + myo-2::GFP],*
>
> *kasEx79 [punc-3_558bp>unc-3 RNAi#n2 + myo-2::GFP],*
>
> *kasEx38 [pcho-1_280bp>lin-39 cDNA OE#1 + myo-2::GFP],*
>
> *kasEx39 [pcho-1_280bp>lin-39 cDNA OE#n2 + myo-2::GFP],*
>
> *kasEx41 [pcho-1_280bp>lin-39 cDNA OE#n9 + myo-2::GFP].*

All reporter strains used in this study are shown in **Table S4**.

### Forward genetic screen

EMS mutagenesis was performed on *unc-3 (n3435); ynIs40 [flp-11::GFP]* animals using standard procedures [67]. Mutagenized L4 animals were visually screened at a dissecting fluorescent microscope for changes in *flp-11::GFP* expression in VNC MNs. One mutant (*kas1*) was retrieved.

### Generation of transgenic reporter animals

Reporter gene fusions for *cis*-regulatory analysis of terminal identity genes were made using either PCR fusion [68] or Gibson Assembly Cloning Kit (NEB #5510S). Targeted DNA fragments were fused (ligated) to *tagrfp* coding sequence, which was followed by *unc-54 3’ UTR.* The TOPO XL PCR cloning kit was used to introduce the PCR fusion fragments into the pCR-XL-TOPO vector (Invitrogen). Mutations on LIN-39 motifs were introduced via mutagenesis PCR. The product DNA fragments were either injected into young adult *pha-1(e2123)* hermaphrodites at 50ng/µl using *pha-1* (pBX plasmid) as co-injection marker (50 ng/µl) and further selected for survival, or injected into young adult N2 hermaphrodites at 50ng/µl (plus 50ng/µl pBX plasmid) using *myo-2::gfp* as co-injection marker (3 ng/µl) and further selected for GFP signal.

The fosmid clone WRM0616aE11 (genomic region: III:7519128..7554793) (Source BioScience) that contains the entire *lin-39* locus was linearized by restriction enzyme digestion, mixed with sonicated bacterial genomic DNA (12 ng/µl) and injected into young adult N2 hermaphrodites at 15 ng/µl using *myo-2::gfp* as co-injection marker (3 ng/µl).

### Generation of transgenic animals for RNAi or over-expression

The cDNA (for over-expression) or the exon-rich genomic region (for RNAi) of *unc-3* and *lin-39* were amplified and then ligated to cholinergic (*cho-1, unc-3*) or GABAergic (*unc-47*) promoters using Gibson Assembly Cloning Kit (NEB #5510S). For *unc-3* RNAi, we targeted exons 2-5 with the following primers: FRW: GTCTGTAAAAGATGAGAACCAGCGG, RVS: CTGTCAATAATAACTGGATCGCTGG. For *lin-39* RNAi, we targeted exons 3-5 with the following primers: FRW: gtggtcaaactccgaacttaaagtg, RVS: gaaggggcgagaaatgtgtgataac. For over-expression constructs, DNA products were purified using a PCR purification protocol (QIAGEN), and then injected into young adult WT hermaphrodites at 50 ng/µl together with 50 ng/µl pBS plasmid (filler DNA) and 3 ng/µl of *myo-2::gfp* (co-injection marker). For RNAi constructs, complementary sense and anti-sense exon-rich genomic regions of *unc-3* and *lin-39* were PCR purified and injected into young adult WT or *unc-3 (n3435)* hermaphrodites each at 100 ng/µl with *myo-2::gfp* as co-injection marker (3 ng/µl) following previously established procedures [69].

### Targeted genome engineering

To generated the *lin-39 (kas9 [lin-39::mNG::AID])* allele, CRISPR/Cas9 genome editing was employed to introduce the *mNG::3xFLAG::AID* cassette into the *lin-39* gene locus before the stop codon. Editing was performed as previously described [70].

### Temporally-controlled protein degradation

AID-tagged proteins are conditionally degraded when exposed to auxin in the presence of TIR1 [33]. Animals carrying auxin-inducible alleles of *lin-39 (kas9 [lin-39::mNG::AID])* or *unc-3 (ot837 [unc-3::mNG::AID])* [47] were crossed with ieSi57 [*eft-3prom::tir1*] animals that express TIR1 ubiquitously. Auxin (indole-3-acetic acid [IAA]) was dissolved in ethanol (EtOH) to prepare 400 mM stock solutions which were stored at 4°C for up to one month. NGM agar plates with fully grown OP50 bacteria were coated with auxin solution to a final concentration of 4 mM, and allowed to dry overnight at room temperature. To induce protein degradation, worms of the experimental strains were transferred onto auxin-coated plates and kept at 20°C. As a control, worms were transferred onto EtOH-coated plates instead. Auxin solutions, auxin-coated plates, and experimental plates were shielded from light.

### Microscopy

Worms were anesthetized using 100mM of sodium azide (NaN_3_) and mounted on a 4% agarose pad on glass slides. Images were taken using an automated fluorescence microscope (Zeiss, Axio Imager.Z2). Acquisition of several z-stack images (each ∼1 µm thick) was taken with Zeiss Axiocam 503 mono using the ZEN software (Version 2.3.69.1000, Blue edition). Representative images are shown following max-projection of 1-8 µm Z-stacks using the maximum intensity projection type. Image reconstruction was performed using Image J software [71].

### Chromatin Immunoprecipitation (ChIP)

ChIP assay was performed as previously described [50, 72] with the following modifications. Synchronized L1 worms (wild-type [N2] and *lin-39 (kas9 [lin-39::mNG::3xFLAG::AID])* strains) were cultured on 10 cm plates seeded with OP50 at 20°C overnight. Early L3 worms were cross-linked and resuspended in FA buffer supplemented with protease inhibitors (150 mM NaCl, 10 µl 0.1 M PMSF, 100 µl 10% SDS, 500 µl 20% N-Lavroyl sarsosine sodium, 2 tablets of cOmplete ULTRA Protease Inhibitor Cocktail [Roche Cat.# 05892970001] in 10ml FA buffer). The sample was then sonicated using a Covaris S220 at the following settings: 200 W Peak Incident Power, 20% Duty Factor, 200 Cycles per Burst for 60 seconds. Samples were transferred to centrifuge tubes and spun at the highest speed for 15 min. The supernatant was transferred to a new tube, and 5% of the material was saved as input and stored at −20°C. Twenty (20) µl of equilibrated anti-FLAG M2 magnetic beads (Sigma-Aldrich M8823) were added to the remainder. Wild-type (N2) worms do not carry the FLAG tag and serve as negative control. The *lin-39 (kas9 [lin-39::mNG::3xFLAG::AID])* CRIPSR-generated allele was used in order to precipitate the immunocomplex comprising the endogenous LIN-39 protein and the bound DNA. The immunocomplex was incubated and rotated overnight at 4°C. On the next day, the beads were washed at 4°C twice with 150 mM NaCl FA buffer (5 min each), once with 1M NaCl FA buffer (5 min). The beads were transferred to a new centrifuge tube and washed twice with 500 mM NaCl FA buffer (10 min each), once with TEL buffer (0.25 M LiCl, 1% NP-40, 1% sodium deoxycholate, 1mM EDTA, 10 mM Tris-HCl, pH 8.0) for 10 min, twice with TE buffer (5 min each). The immunocomplex was then eluted in 200 µl elution buffer (1% SDS in TE with 250 mM NaCl) by incubating at 65°C for 20 min. The saved input samples were thawed and treated with the ChIP samples as follows. One (1) µl of 20 mg/ml proteinase K was added to each sample and the samples were incubated at 55°C for 2 hours and then at 65°C overnight (12-20 hours) to reverse cross-link. The immunoprecipitated DNA was purified with Ampure XP beads (A63881) according to manufacturer’s instructions.

### Real-time quantitative PCR (qPCR) analysis of ChIP DNA

qPCR analysis of ChIP DNA was performed to probe enrichment of predicted LIN-39 binding sites at target genes. Two biological replicates were included. The primers used are provided in 5’-3’ orientation: *acr-2* (FRW: acattcgcaccaacaaagcg; RVS: aaaggacggacccaacagac), *del-1* (FRW: tacagttataggtaccattgtcgag; RVS: tttacagagaaaatttgaattgccg), *nca-1* (FRW: gcatccgatggaaagggtcc; RVS: ttcccttgtctagcagtcgg), *slo-2* (FRW: caagtttcactgggcgacaaa; RVS: agggaactcaaggcttacatgg), *unc-129* (FRW: attcgtgtctcgcagggaac; RVS: atagaggaaccggcaaaggtg), *oig-1 site #4* (FRW: atcacatcatttcaaaggagagcc; RVS: tgagtaatgaggcgtaacacgg), *oig-1 site #5* (FRW: acgtgtttcttgtgcttccg; RVS: caatgacggaaataaagattgtgcg), *oig-1 site #14* (FRW: ccaaactctataatcccaaccccc; RVS: tcgaaacgtcgaaaaatgggag). The amplification was conducted in a QuantStudio 3 using the Power SYBR Green PCR Master Mix (ThermoFisher Cat.# 4367659), with the following program: Step 1: 95°C for 10 min; Step 2: 95°C for 15 s; Step 3: 60°C for 1 min. Repeat steps 2-3 for 40 times.

### Motor neuron identification

Motor neuron (MN) subtypes were identified based on combinations of the following factors: (i) co-localization with fluorescent markers with known expression pattern, [32] invariant cell body position along the ventral nerve cord, or relative to other MN subtypes, (iii) MN birth order, and (iv) number of MNs that belong to each subtype.

### Bioinformatic analysis

To predict the UNC-3 binding site (COE motif) in the *cis*-regulatory region of *unc-129, del-1, acr-2, nca-1* and *slo-2*, we used the MatInspector program from Genomatix (http://www.genomatix.de) (Cartharius, 2005 #3476). The Position Weight Matrix (PWM) for the LIN-39 binding site is catalogued in the CIS-BP (Catalog of Inferred Sequence Binding Preferences) database (Weirauch, 2014 #8691) (http://cisbp.ccbr.utoronto.ca/). To identify putative LIN-39 sites on the *cis*-regulatory regions of *unc-129, del-1, acr-2, nca-1, slo-2, oig-1, ser-2, flp-11 and lin-39,* we used FIMO (Find Individual Motif Occurrences; (Grant, 2011 #8201)), which is one of the motif-based sequence analysis tools of the MEME (Multiple Expectation maximization for Motif Elicitation) bioinformatics suite (http://meme-suite.org/). To predict the binding site for the transcription factor UNC-30, we performed FIMO analysis using the UNC-30 binding motif (WNTAATCHH) described in [36]. The p-value threshold for the analysis was set at p < 0.001 for *del-1, nca-1, slo-2, ser-2, flp-11* and *lin-39,* and at p < 0.01 for *unc-129, acr-2 and oig-1*.

### Automated worm tracking

Worms were maintained as mixed stage populations by chunking on NGM plates with *E. coli* OP50 as the food source. The day before tracking, 30-40 L4 larvae were transferred to a seeded NGM plate and incubated at 20°C for approximately 24 hours. Five adults are picked from the incubated plates to each of the imaging plates (see below) and allowed to habituate for 30 minutes before recording for 15 minutes. Imaging plates are 35 mm plates with 3.5 mL of low-peptone (0.013% Difco Bacto) NGM agar (2% Bio/Agar, BioGene) to limit bacteria growth. Plates are stored at 4°C for at least two days before use. Imaging plates are seeded with 50µl of a 1:10 dilution of OP50 in M9 the day before tracking and left to dry overnight with the lid on at room temperature.

### Behavioral feature extraction and analysis

All videos were analyzed using Tierpsy Tracker [73] to extract each worm’s position and posture over time. These postural data were then converted into a set of behavioral features as previously described [53]. From the total set of features, we only considered 48 that are related to midbody posture and motion, as well as the midbody width (see **Table S3** for feature descriptions and their average values for each strain). For each strain comparison, we performed unpaired two-sample t-tests independently for each feature. The false discovery rate was controlled at 5% across all strain and feature comparisons using the Benjamini Yekutieli procedure [74]. The p-value threshold to control the false discovery rate at 0.05 is 0.0032.

### Cloning, western blot, and immunoprecipitation

UNC-3, and LIN-39 cDNAs were cloned into the mammalian expression vectors pcDNA 3.1(+)-C-Flag plasmid and the pcDNA 3.1(+)-N-Myc plasmid by GeneScript, to generate C-terminus Flag-tagged UNC-3 and N-terminus Myc-tagged LIN-39. The constructs were verified by sequencing at the sequencing core facility of University of Chicago. The tagged proteins were expressed in HEK293 cells. Protein expression was detected by standard western blot. Expression of Myc tagged LIN-39 was detected using anti-Myc (Abcam, #ab9106), expression of Flag-tagged UNC-3 in the total cell lysate was detected using mouse anti-Flag (Sigma, #F3165), expression of Flag-tagged UNC-3 in the IP was detected using rabbit anti-Flag (Sigma, #SAB4301135). Immunoprecipitation of Flag-tagged UNC-3 was performed using Flag antibody coated beads (Sigma, #A2220). For the IP, the “Clean-Blot IP Detection Reagent” (Thermo Fisher, #21230) was used as secondary antibody.

### Statistical analysis

For data quantification, graphs show values expressed as mean ± standard deviation (STDV). The statistical analyses were performed using the unpaired *t*-test (two-tailed). Calculations were performed using the Evan’s Awesome A/B Tools online software (https://www.evanmiller.org/ab-testing/t-test.html). Differences with *p* < 0.05 were considered significant.

## SUPPLEMENTARY FIGURE LEGENDS

**Supplementary Figure 1:**
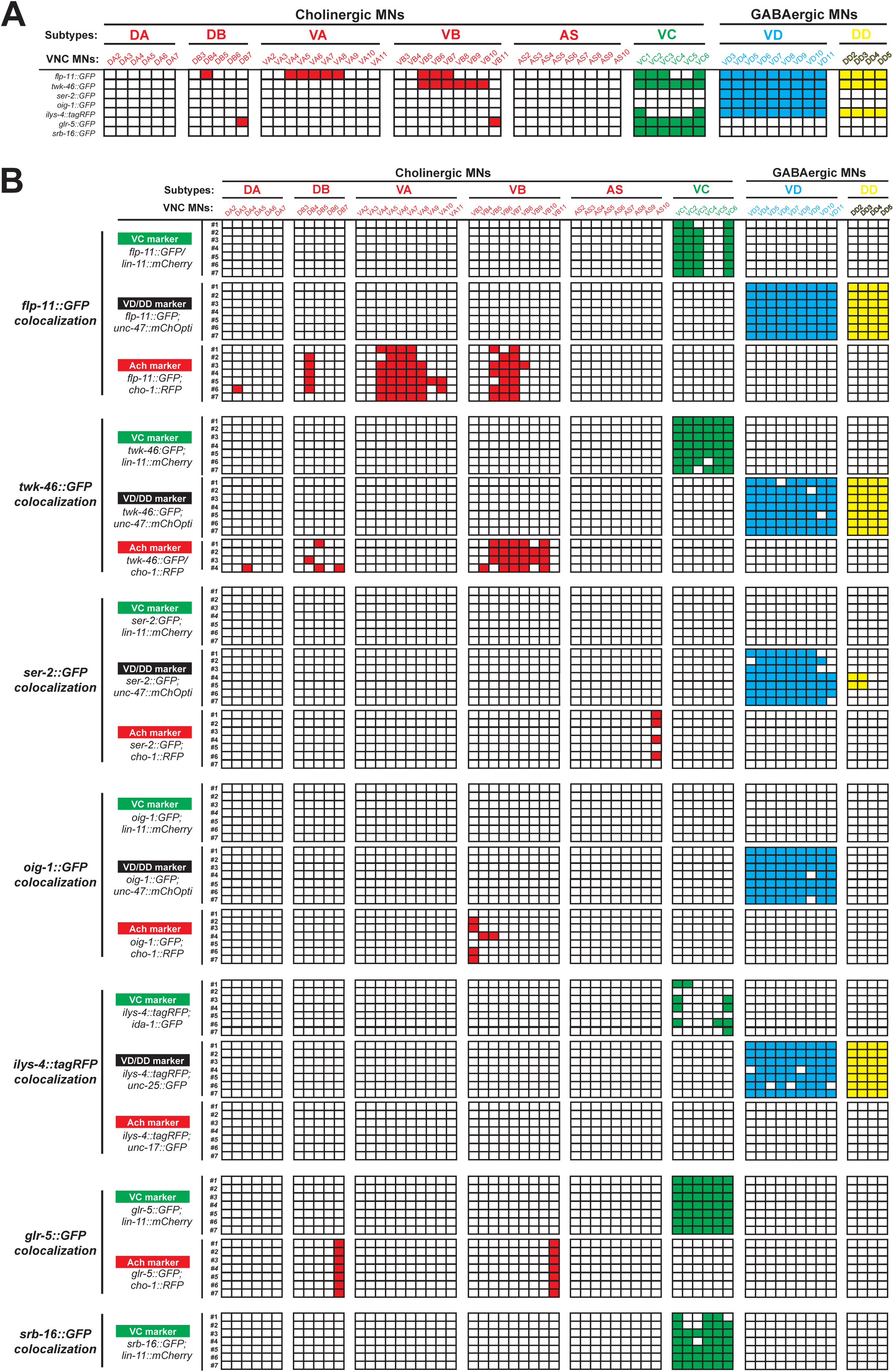
Detailed characterization of the expression pattern of VC and VD terminal identity markers. **A**: An expression map of VC and VD terminal identity genes with single-cell resolution at larval stage 4 (L4). Each column represents an individual motor neuron (MN) in the ventral nerve cord (VNC) of a hermaphrodite worm. MN subtypes are color-coded. This table represents a summary of the co-localization analysis described in panel B. Apart from the 7 genes listed, we also examined 5 additional terminal identity genes (bra-1, dhc-1, rgs-4, vhp-1, vps-25) but found no evidence for expression in VD or VC neurons. **B**: Co-localization analysis of VD and VC terminal identity genes. Each column represents an individual MN in the VNC of a hermaphrodite worm. Each row represents a randomly-selected worm co-expressing the respective VD or VC marker and one of the known identity markers: *unc-47::mChOpti or unc-25::gfp* for GABAergic (VD and DD) MN identity, *cho-1::rfp or unc-17::gfp* for cholinergic MN identity and *lin-11::mCherry or ida-1::gfp* for VC MN identity. A color-filled lattice indicates co-expression of the known identity marker and the selected marker in the individual MN.

**Supplementary Figure 2:**
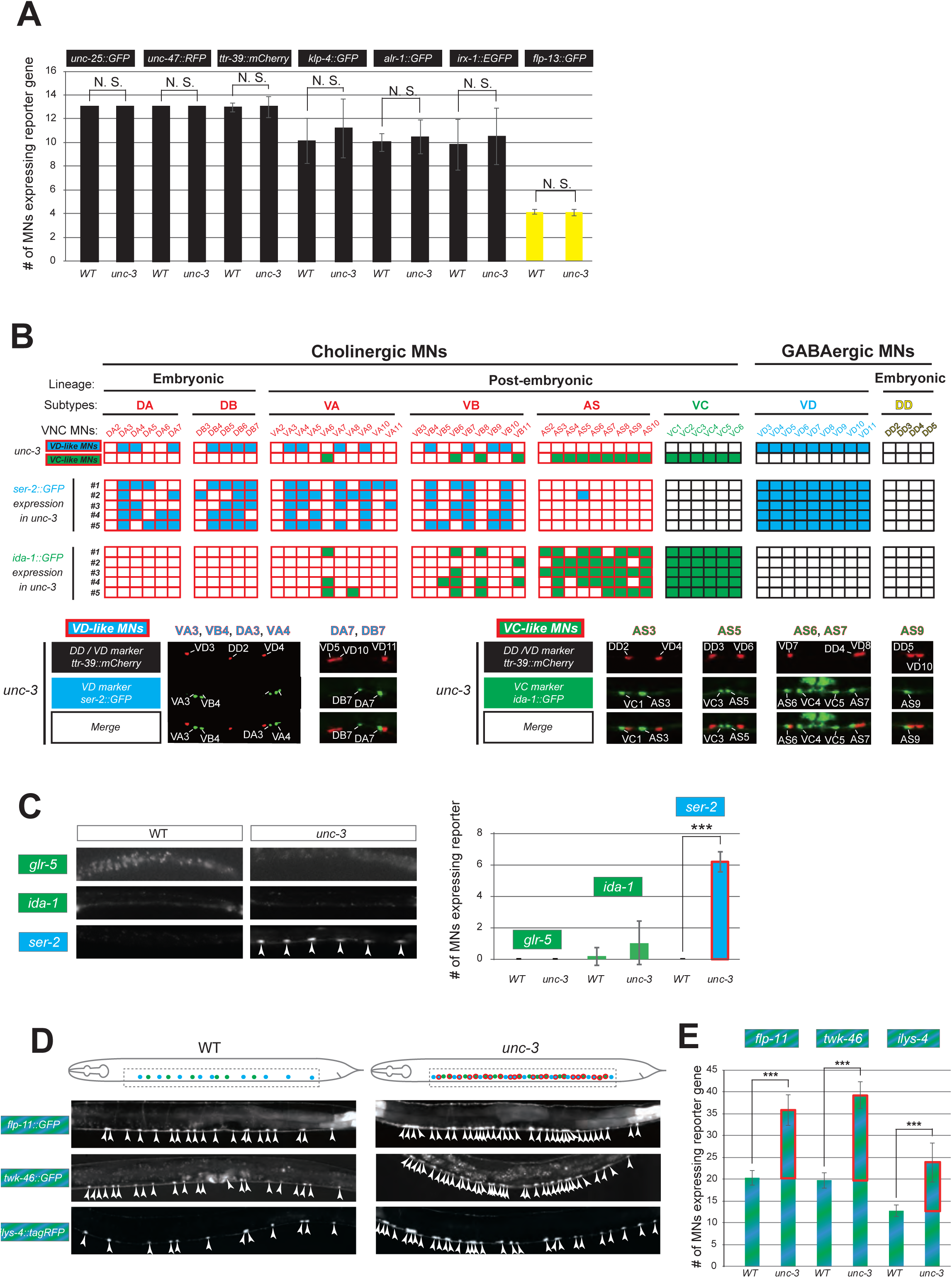
UNC-3 selectively prevents expression of VD and VC terminal identity features in distinct cholinergic MNs. **A:** Quantification of six VD/DD (*unc-25::gfp, unc-47::rfp, ttr-39::mCherry, klp-4::gfp, alr-1::gfp and irx-1::egfp*) and one DD (*flp-13::gfp*) markers in WT and *unc-3 (n3435)* animals at L4. No statistically significant effects were observed in *unc-3* mutants. N > 13. N. S: not significant. **B**: A table showing that VD and VC identity genes are ectopically expressed in distinct cholinergic MNs upon *unc-3* depletion. Top two rows: A summary map of VD and VC gene expression with single-cell resolution in *unc-3* mutants, which is based on the co-localization analysis shown below. Each column represents an individual MN in the VNC of an *unc-3*-depleted hermaphrodite worm. Five randomly selected worms that co-express VD/DD marker *ttr-39::mCherry* and VD marker *ser-2::gfp* or VC marker *ida-1::gfp* are analyzed. The expression of *ttr-39::mCherry* is not affected by *unc-3* depletion, which serves as a positional landmark for reference. This landmark together with the invariant MN cell body position along the VNC enable the identification of distinct *unc-3*-depleted MNs that lose cholinergic features and concomitantly gain either “VD-like” or “VC-like” features. Below this table, examples of the co-localization analysis are provided. Representative magnified images are shown. The identity of *unc-3*-depleted MNs that acquire VC or VD terminal features is shown. Note that VC-like terminal features are mainly acquired by MNs of the AS cholinergic subtype, whereas VD-like terminal features are acquired by select members of four cholinergic MN subtypes (DA, DB, VA, VB). **C**: The DA and DB cholinergic MNs are born embryonically and therefore present in the VNC at L1. Corroborating our results from panel B, only the VD terminal identity marker (*ser-2*) shows ectopic expression in DA and DB neurons of unc-3 mutants at L1. This is not the case for the VC terminal identity markers (*glr-5, ida-1*). Representative images are shown on the left. Arrowheads point to MN cell bodies with ectopic gfp marker expression. Green fluorescence signal is shown in white for better contrast. Quantification is on the right. Ectopic expression in *unc-3*-depleted MNs is highlighted with a red rectangle. N > 22. *** p < 0.001. **D:** Terminal identity markers of VD/VC MNs (*flp-11, twk-46 and ilys-4*) are ectopically expressed in *unc-3*-depleted cholinergic MNs. Representative images of larval stage 4 (L4) hermaphrodites are shown. Similar results were obtained in adult animals. Arrowheads point to MN cell bodies with marker expression. Fluorescence signal is shown in white for better contrast. Dotted black box indicates imaged area. **E**: Quantification of data shown in panel D. Ectopic expression is highlighted with a red rectangle. N > 18. *** p < 0.001.

**Supplementary Figure 3:**
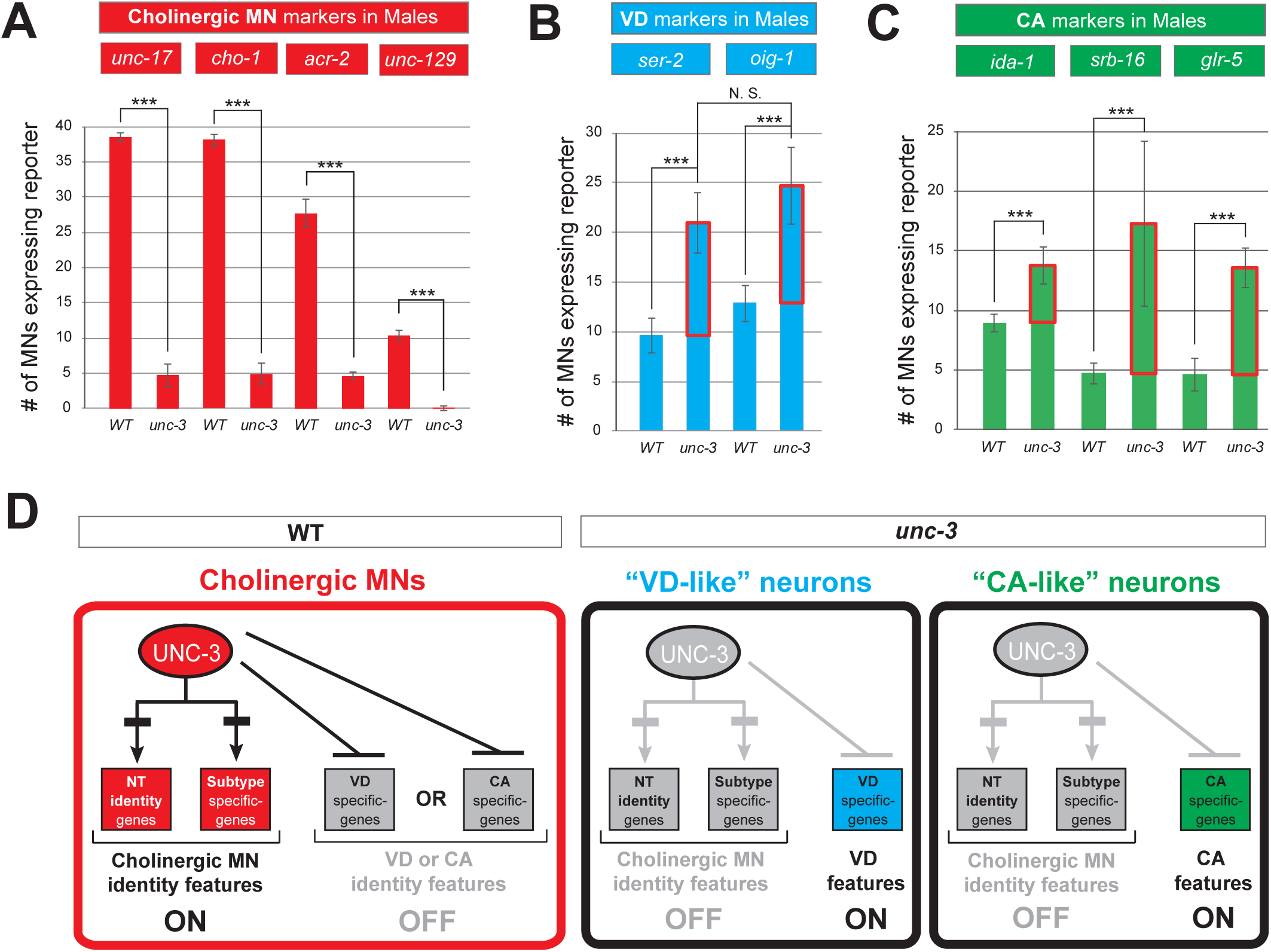
The dual role of UNC-3 in cholinergic MNs extends to both *C. elegans* sexes. **A:** Quantification of four sex-shared cholinergic MN markers (*unc-17, cho-1, acr-2*, *unc-129*) expression in WT and *unc-3 (n3435)* male animals at L4. N > 13. *** p < 0.001. **B**: Quantification of VD marker (*ser-2 and oig-1*) expression in L4 stage WT and *unc-3 (n3435)* male animals. Ectopic expression is highlighted with a red rectangle. No significant difference was found between the number of MNs ectopically expressing each marker. N > 11. *** p < 0.001. N. S: not significant. **C:** Quantification of the male-specific CA markers (*ida-1, srb-16 and glr-5*) in young adult stage WT or *unc-3 (n3435)* male animals. Ectopic expression is highlighted with a red rectangle. N > 14. *** p < 0.001. **D:** Schematic summarizing the dual role of UNC-3 in *C. elegans* males.

**Supplementary Figure 4:**
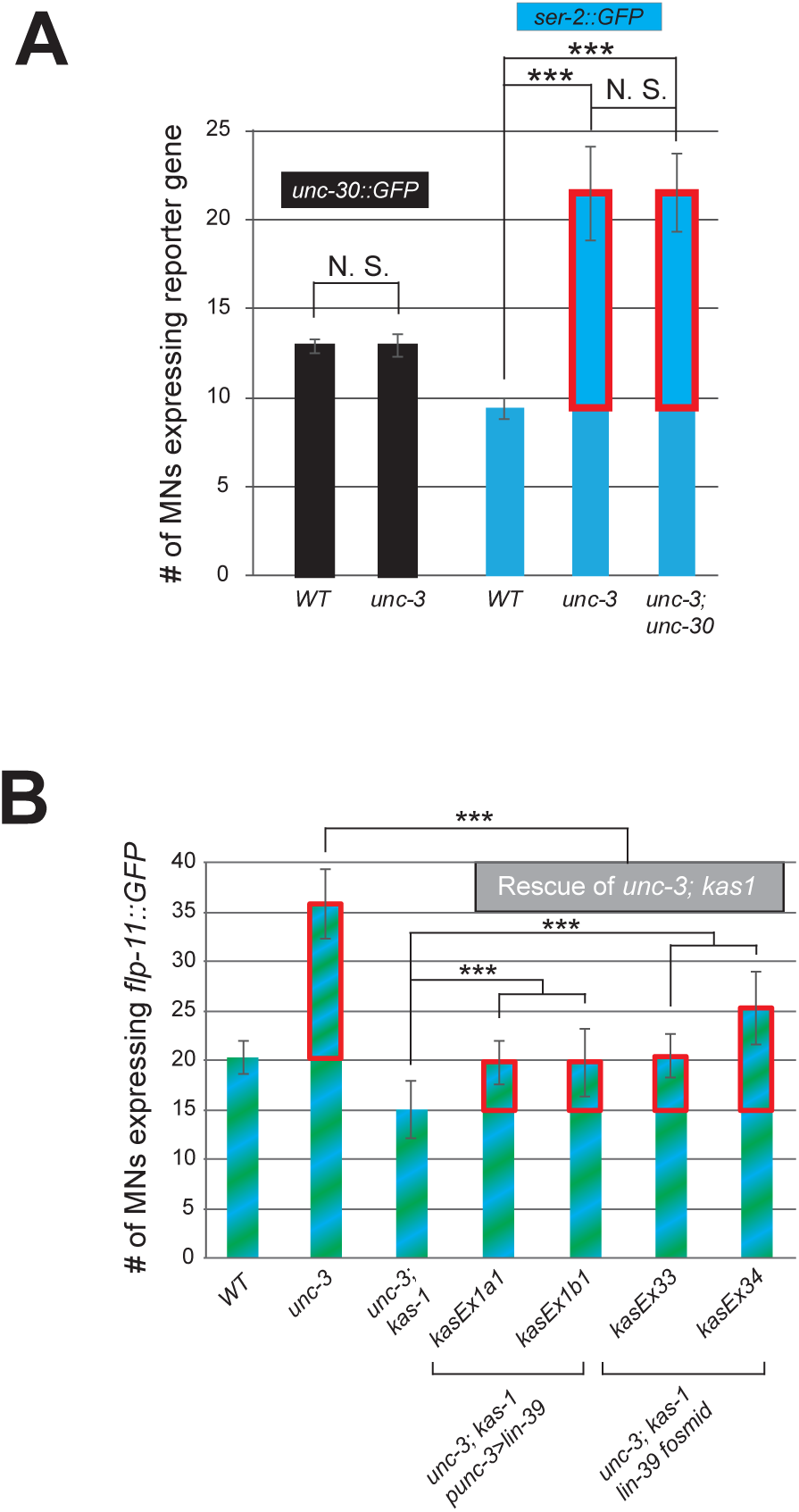
Ectopic expression of VD terminal identity markers in *unc-3* mutants requires LIN-39 but not UNC-30. **A**: Quantification of *unc-30::gfp* and VD marker *ser-2::gfp* in WT and *unc-3 (n3435)* animals at L4, with *ser-2::gfp* further examined in *unc-3 (n3435); unc-30 (e191)* double mutants. Expression of *ser-2::gfp* is equally affected in *unc-3 (n3435)* single and *unc-3 (n3435); unc-30 (e191)* double mutants. N > 15. *** p < 0.001. N. S: not significant. **B**: Quantification of the VD/VC marker *flp-11::gfp* expression in L4 stage WT, *unc-3 (n3435)* mutant, *unc-3 (n3435); kas1*, and transgenic animals that rescue *unc-3 (n3435); kas1* background with expression of *lin-39* in cholinergic MNs (*Punc-3 > lin-39*) or expression of GFP-tagged *lin-39* fosmid. Ectopic expression is highlighted with a red rectangle. Two independent transgenic lines were used for *Punc-3 > lin-39 OE* and *lin-39* fosmid. Of note, the partial rescue observed with the *Punc-3 > lin-39 OE* lines likely arises because the fragment of *unc-3* promoter used to drive *lin-*39 is not expressed in all 39 cholinergic MNs. N > 16. *** p < 0.001.

**Supplementary Figure 5:**
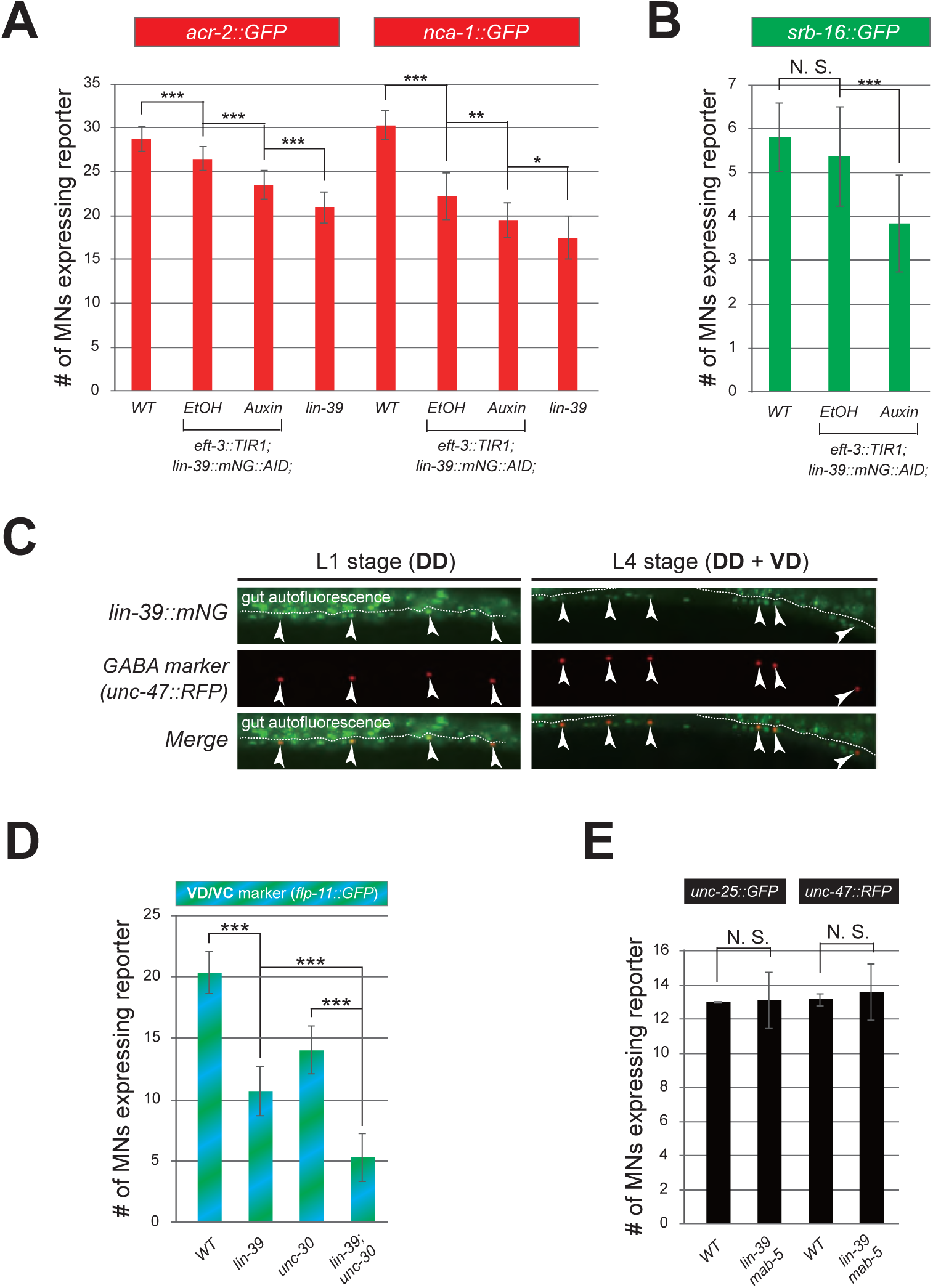
LIN-39 is continuously required to activate distinct terminal identity genes in sex-shared and sex-specific cholinergic MNs. **A:** Auxin or ethanol (control) were administered at larval stage 4 (L4) on *lin-39::mNG::3xFLAG::AID; eft-3::TIR1* animals carrying the sex-shared cholinergic MN markers *acr-2::gfp* and *nca-1::gfp*. Images were taken at the young adult stage (day 1 for *acr-2::gfp* and day 2.4 for *nca-1::gfp*) and the number of MNs expressing these markers was quantified. A statistically significant decrease is evident in the auxin-treated animals compared to EtOH-treated controls. For comparison, quantification of marker expression is also provided in WT and *lin-39 (n1760)* animals. N > 12. * p < 0.05, ** p < 0.01, *** p < 0.001. **B:** Auxin or ethanol (control) were administered at larval stage 3 (L3) on *lin-39::mNG::3xFLAG::AID; eft-3::TIR1* animals carrying the VC marker *srb-16::gfp.* Quantification was performed at the young adult stage (day 1.5). A significant decrease in the number of MNs expressing the VC marker was evident in the auxin-treated animals compared to EtOH-treated controls. For comparison, quantification of marker expression is also provided in WT animals. N > 15. ** p < 0.01, *** p < 0.001. N. S: not significant. **C:** Representative images of L1- and L4-stage animals co-expressing *unc-47::rfp* (labels only DD at L1, labels both DD and VD at L4) and endogenous *lin-39* marker (*lin-39::mNG*). Arrowheads point to cell bodies of DD and VD MNs co-expressing both markers. White dotted line indicates the boundary of intestinal tissue (gut), which tends to be autofluorescent in the green channel. **D:** Quantification of VD/VC marker *flp-11::gfp* in WT, *lin-39 (n1760), unc-30 (e191)* and *lin-39 (n1760); unc-30 (e191)* double mutant animals at L4 stage. Double mutants showed a more severe reduction in *flp-11::gfp* expression compared to each single mutant. N > 20. *** p < 0.001. **E:** Expression of GABA biosynthesis markers (*unc-25, unc-47*) is not affected in *lin-39 (n1760); mab-5 (e1239)* double mutants at the L4 stage. N > 15; N. S: not significant.

**Supplementary Figure 6:**
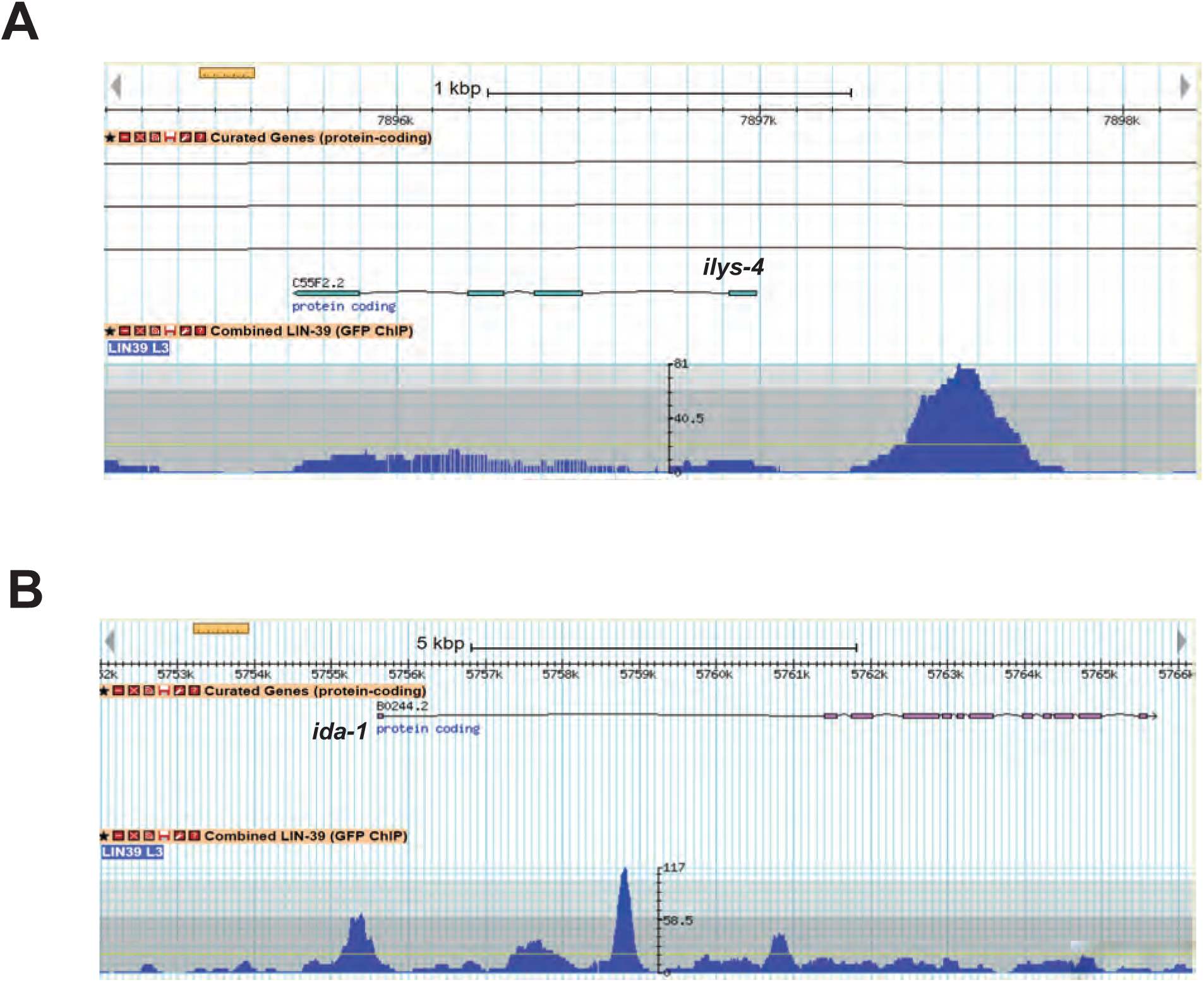
LIN-39 binds directly to the *cis*-regulatory region of VD and VC terminal identity genes. Genome browser view of *ilys-4* (**A**) and *ida-1* (**B**) loci (www.wormbase.org). ChIP-seq for LIN-39 shows that it binds directly to the *cis*-regulatory region of VD and/or VC terminal identity genes (*ilys-4, ida-1*). Data were generated through the modENCODE project and deposited into WormBase, from which these images were downloaded.

**Supplementary Figure 7:**
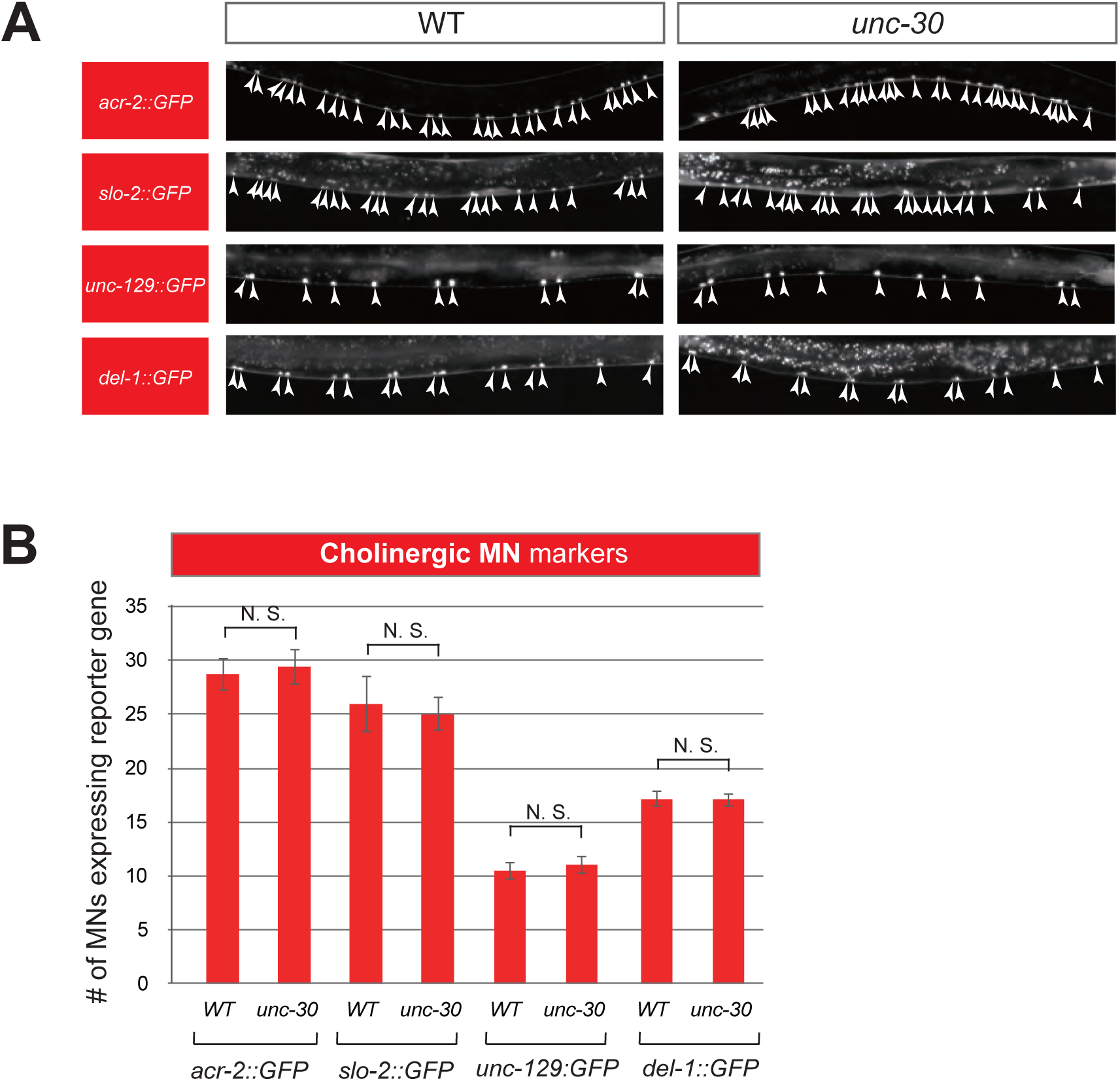
UNC-30 appears not to have a dual role in GABAergic MNs. **A**: Several terminal identity markers of cholinergic neurons (*acr-2, slo-2, unc-129, del-1*) are not ectopically expressed in *unc-30*-depleted GABAergic MNs. A strong loss-of-function allele *e191* for *unc-30* was used [27, 37]. Representative images of larval stage 4 (L4) hermaphrodites are shown on the left. Arrowheads point to MN cell bodies with *gfp* marker expression. Green fluorescence signal is shown in white for better contrast. **B**: Quantification of data presented in panel A. N. S: not significant.

**Supplementary Figure 8:**
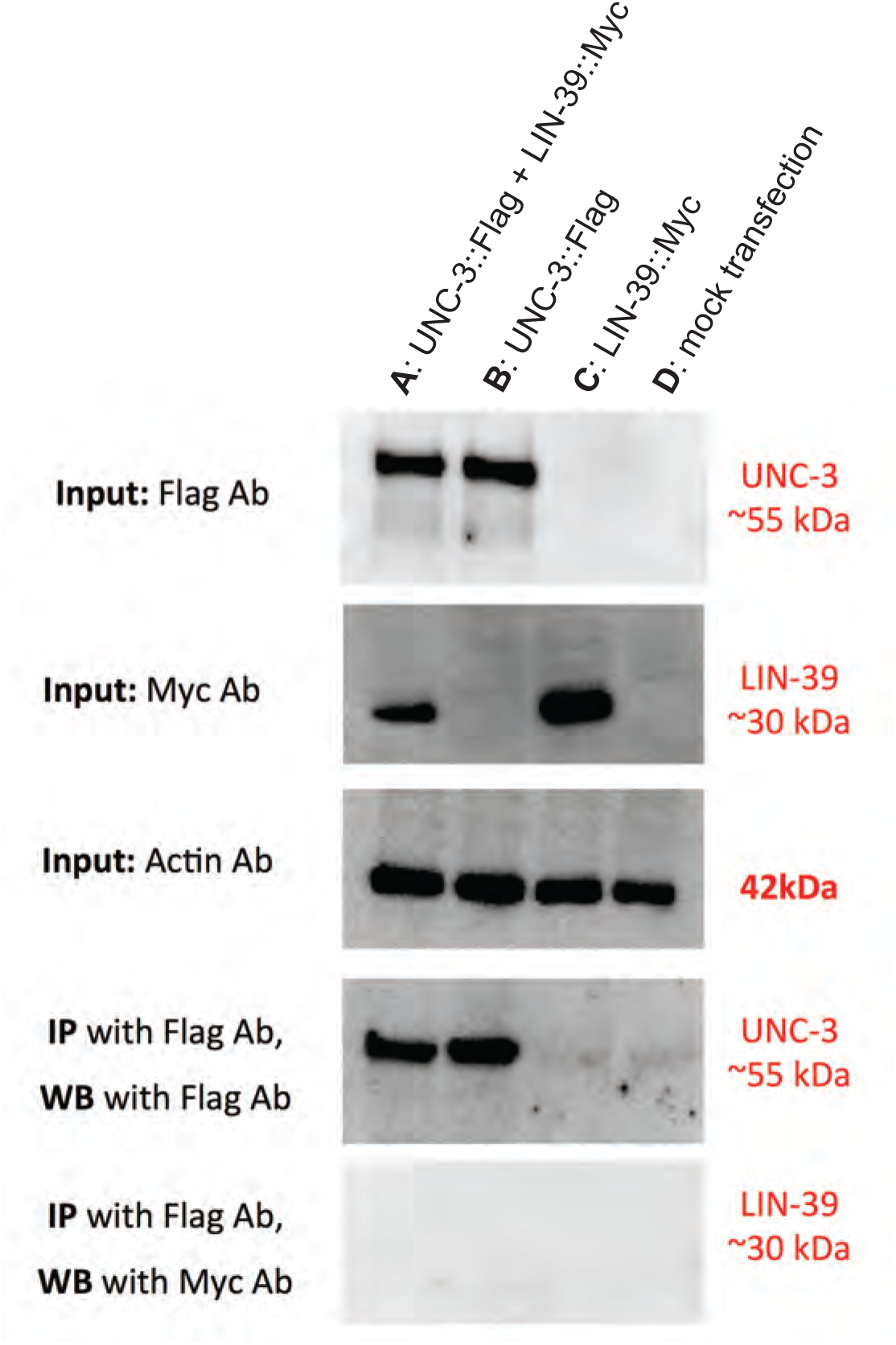
UNC-3 does not physically interact with LIN-39 in a heterologous system. (A) HEK293 cells were transfected with two mammalian expression plasmids encoding Flag-tagged UNC-3 and a plasmid encoding Myc-tagged LIN-39, or (B) with a single plasmid encoding the Flag-tagged UNC-3, or (C) with a single plasmid encoding the Myc-tagged LIN-39, or (D) were mock transfected. 10 ug of the protein lysate were subjected to western blot analysis using antibodies against Flag, Myc and Actin (upper three panels). 100 ug of these cell lysates were used for immunoprecipitation of the Flag-tagged UNC-3, using anti-Flag antibody coated beads (lower two panels).

**Supplementary Figure 9:**
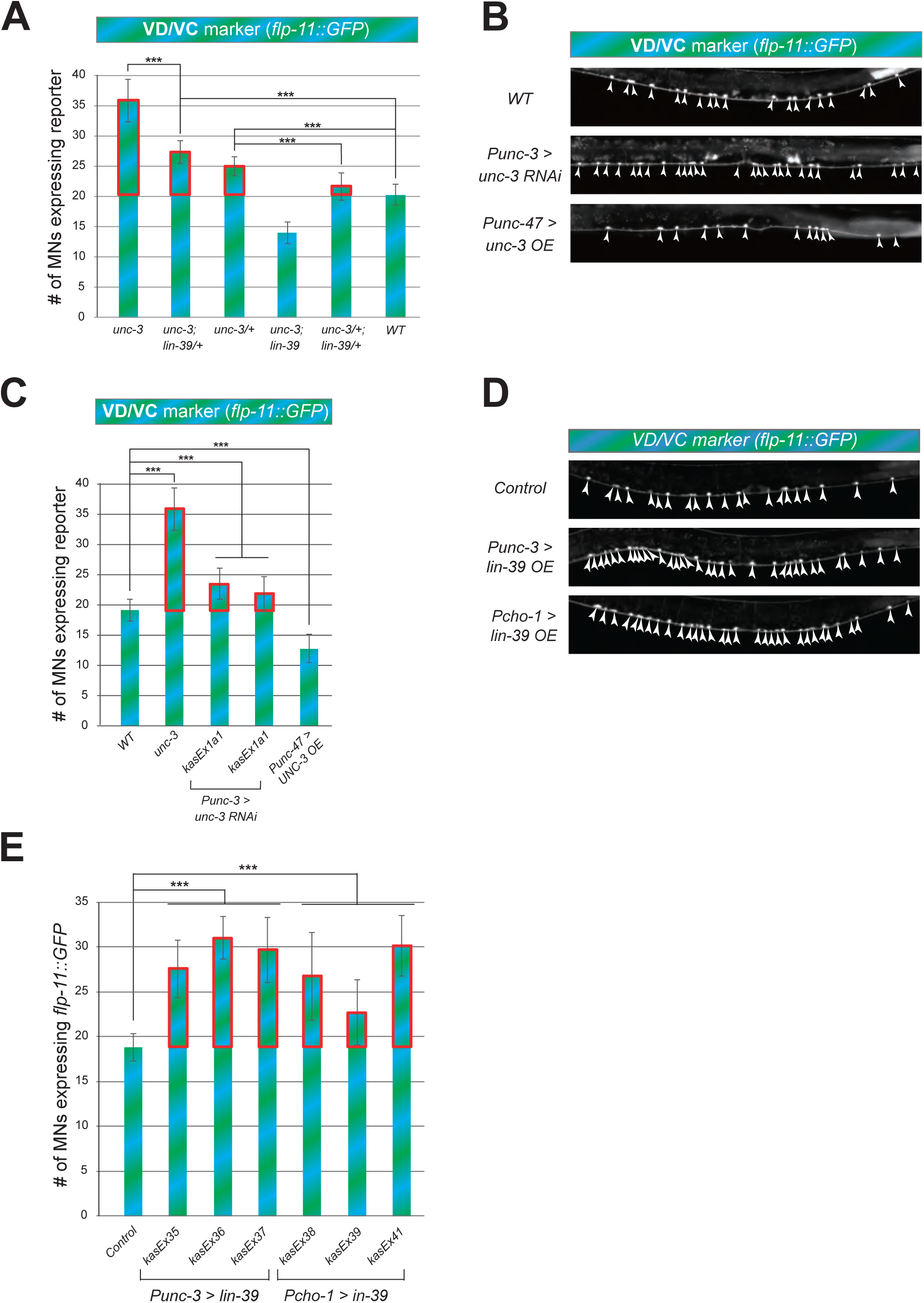
UNC-3 and LIN-39 levels are crucial for ectopic expression of VD/VC terminal identity marker *flp-11*. **A:** Quantification of the VD/VC marker *flp-11::gfp* in *unc-3 (n3435), unc-3 (n3435); lin-39 (n1760)/+, unc-3 (n3435)/+, unc-3 (n3435); lin-39 (n1760), unc-3 (n3435)/+; lin-39 (n1760)/+*, and WT animals at L4. Ectopic expression is highlighted with a red rectangle. N > 18. *** p < 0.001. **B**: Representative images of the VD/VC marker *flp-11::gfp* expression in L4 stage WT or transgenic animals that either down-regulate *unc-3* in cholinergic MNs (*Punc-3 > unc-3 RNAi*) or over-express *unc-3* in GABAergic (including VD) neurons (*Punc-47 > unc-3 OE*). Arrowheads point to MN cell bodies with *gfp* expression. Green fluorescence signal is shown in white for better contrast. **C:** Quantification of data shown in panel B. Ectopic expression is highlighted with a red rectangle. Two independent transgenic lines were used for *Punc-3 > unc-3 RNAi*. N > 17. *** p < 0.001. Quantification of *flp-11::gfp* is also shown in WT and *unc-3 (n3435)* null animals for comparison. **D**: Representative images of the VD/VC marker *flp-11::gfp* expression in L4 stage WT or transgenic animals that over-express *lin-39* in cholinergic MNs driven by two different cholinergic MN promoters(*Punc-3 > lin-39 OE, Pcho-1 > lin-39 OE*). Arrowheads point to MN cell bodies with marker expression. Green fluorescence signal is shown in white for better contrast. **E:** Quantification of data shown in panel D. Ectopic expression is highlighted with a red rectangle. Three independent transgenic lines were used for *Punc-3 > lin-39 OE* and *Pcho-1 > lin-39 OE*. N =19. *** p < 0.001.

**Supplementary Figure 10:**
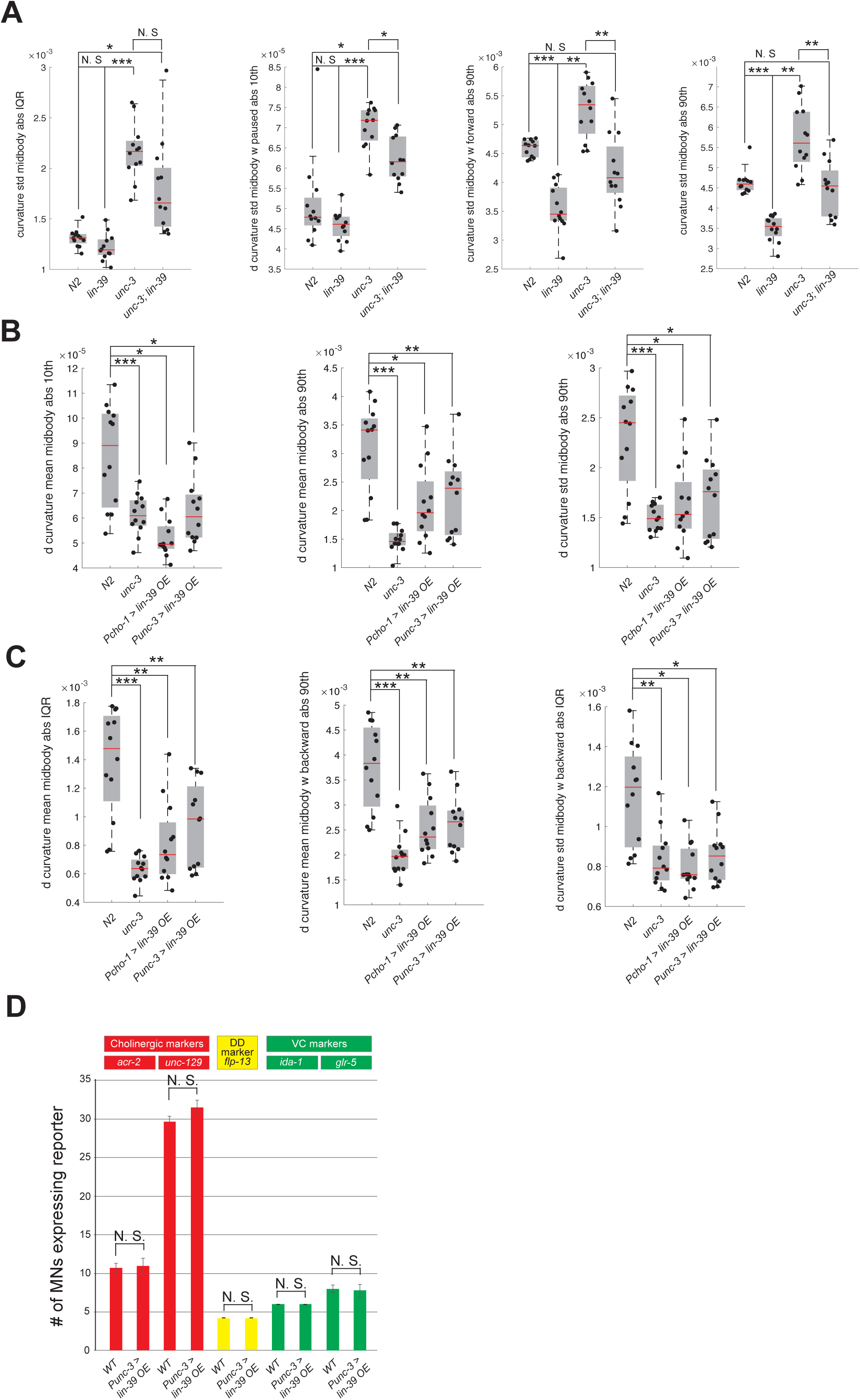
Automated worm tracking analysis on *unc-3* and *unc-3; lin-39* mutants. **A**: Additional mid-body locomotion features that are significantly affected in *unc-3 (n3435)* animals, but markedly improved in *unc-3 (n3435); lin-39 (n1760)* double mutant animals. Each black dot represents a single adult animal. The unit for the first two graphs is 1/microns. The unit for the graph on the right is 1/(microns*seconds). N = 12. For a comprehensive list of mid-body features see **Suppl. Table 3**. * p < 0.01, ** p < 0.001, *** p < 0.0001. **B-C**: Additional mid-body locomotion features affected in *unc-3 (n3435)* mutants and animals over-expressing *lin-39* in cholinergic MNs. Each black dot represents a single adult animal. The unit for the y axis is 1/(microns*seconds). N = 12. For a comprehensive list of mid-body features see **Suppl. Table 3**. * p < 0.01, ** p < 0.001, *** p < 0.0001. **D:** Over-expression of LIN-39 in cholinergic MNs (*Punc-3 > LIN-39 OE*) does not affect expression of terminal identity genes specific to cholinergic (*acr-2, unc-129*), GABAergic DD (*flp-13*) or VC (*ida-1, glr-5*) motor neurons. L4 stage animals were imaged and quantified. N > 15.

