## Supplemental Table 1 for "A terminal selector prevents a Hox transcriptional switch to safeguard motor neuron identity throughout life"

**Table S1: UNC-3 binding sites (COE motifs) are not found in the *cis*-regulatory region of VD- and VC-expressed terminal identity genes.**

| Terminal identity gene | Expression | Effect in *unc-3 (-)* | COE motif |
|---|---|---|---|
| *unc-17* | Cholinergic MNs | Loss of expression in MNs | Yes |
| *cho-1* | Cholinergic MNs | | Yes |
| *acr-2* | Cholinergic MNs | | Yes |
| *del-1* | Cholinergic MNs | | Yes |
| *unc-129* | Cholinergic MNs | | Yes |
| *nca-1* | Cholinergic MNs | | Yes |
| *slo-2* | Cholinergic MNs | | Yes |
| *ser-2* | VD | Ectopic expression in MNs | No |
| *oig-1* | VD | | No |
| *flp-11* | DD/VD/VC | | No |
| *twk-46* | DD/VD/VC | | No |
| *ilys-4* | DD/VD/VC | | No |
| *glr-5* | VC | | No |
| *ida-1* | VC | | No |
| *srb-16* | VC | | No |
