## Supplemental Table 2 for "A terminal selector prevents a Hox transcriptional switch to safeguard motor neuron identity throughout life"

**Table S2. Novel LIN-39/Hox targets in cholinergic and GABAergic (VD) motor neurons.**

| LIN-39 targets in cholinergic MNs | | | | | LIN-39 targets in GABAergic VD MNs | | | |
|---|---|---|---|---|---|---|---|---|
| | **COE motif** (relative to ATG) | **LIN-39 sites** (relative to ATG) | **LIN-39** ChIP-qPCR | | | **UNC-30** sites (relative to ATG) | **LIN-39** sites (relative to ATG) | **LIN-39** ChIP-qPCR |
| *unc-129* | *COE1 (-262-240)<br>COE2 (-346-324)<br>COE3 (-458-436)* | *LIN-39 sites:<br>#1 (-275-268)<br>#2 (-321-314)<br>#3 (-376-369)<br>#4 (-444-437)<br>#5 (-616-609)<br>#6 (-819-812)* | *Yes<br>sites #1,#2, #3* | | *oig-1** | *UNC-30 sites:<br>#1 (-423-418)<br>#2 (-955-951)<br>#3 (-1006-1001)<br>#4 (-1181-1176)<br>#5 (-1266-1261)* | *LIN-39 sites:<br>#1 (-172-165)<br>#2 (-406-399)<br>#3 (-728-721)<br>#4 (-843-836)<br>#5 (-974-967)<br>#6 (-1038-1031)<br>#7 (-1126-1119)<br>#8 (-1594-1587)* | *Yes<br>site#5* |
| *del-1* | *COE1 (-132-110)<br>COE2 (-962-941)* | *LIN-39 sites:<br>#1 (-288-281)<br>#2 (-416-409)<br>#3 (-1082-1075)<br>#4 (-1146-1135)* | *Yes<br>site#1* | | *ser-2** | *UNC-30 site in pser-2:<br>#1 (-253-246)* | *LIN-39 site in pser-2:<br>#1(+1261+1268)* | *N. D* |
| *acr-2** | *COE2 (-483-461)<br>COE1 (-147-126)* | *LIN-39 sites:<br>#1 (-486-477)<br>#2 (-512-505)<br>#3 (-580-564)<br>#4 (-617-610)<br>#5 (-646-639)<br>#6 (-759-752)<br>#7 (-791-784)* | *Yes<br>site#1* | | *flp-11** | *UNC-30 sites:<br>#1 (-1635-1627)<br>#2 (-2273-2265)* | *LIN-39 site in pflp-11f2b:<br>#1 (-1307-1300)<br>#2 (-1140-1133)<br>#3 (-1069-1062)<br>#4 (-994-987)* | *N. D* |
| *nca-1** | *COE1 (-3935-3913)<br>COE2 (-3291-3269)* | *LIN-39 sites:<br>#1 (-9634-9627)<br>#2 (-9627-9620)<br>#3 (-8852-8845)<br>#4 (-3524-3517)<br>#5 (-1546-1539)* | *Yes<br>site#2* | | | | | |
| *slo-2** | *COE1 (-176-157)<br>COE2 (-217-198)<br>COE3 (-419-399)* | *LIN-39 site #1<br>(-198-190)* | *Yes<br>site#1* | | | | | |

Asterisk (*) highlights novel LIN-39 targets; N. D: Not Determined.

Note 1: All UNC-3 binding sites (COE motifs 22bp) have been previously described in Kratsios et al. 2012.

Note 2: UNC-30 site#2 on *oig-1* was experimentally validated in Howell et al., 2015.
