## Supplemental Table 4 for "A terminal selector prevents a Hox transcriptional switch to safeguard motor neuron identity throughout life"

### Table S4: Reporter strains used in this study

| Strain name | Genotype |
|---|---|
| OH2246 | *otIs107 [ser-2::GFP + lin-15(+)] I* |
| QW192 | *zfIs8 [ser-2::RFP] IV* |
| OH3955 | *pha-1 (e2123); otEx193 [oig-1::GFP]* |
| BL5717 | *inIs179 [ida-1::GFP] II; him-8(e1489) IV* |
| BC14820 | *dpy-5(e907) I; sEx14820 [srb-16::GFP + pCeh361]* |
| AL270 | *icIs270 [glr-5::GFP + lin-15(+)] X* |
| CA1200 | *unc-119(ed3); ieSi57 [eft-3p::TIR1::mRuby::unc-54 3'UTR + Cbr-unc-119(+)] II* |
| NY2040 | *ynIs40 [flp-11p::GFP] V* |
| BC15028 | *dpy-5 (e907) I; sEx15028 [nca-1::GFP]* |
| BW1935 | *unc-119(ed3) III; ctIs43 [dbl-1::GFP + unc-119(+)] V; him-5(e1490)V* |
| CZ631 | *juIs14 [acr-2p::GFP + lin-15(+)] IV* |
| BC10749 | *dpy-5 (e907) I; sEx10749 [slo-2::GFP + pCeh361]* |
| LX929 | *vsIs48 [unc-17::GFP] X* |
| OH13646 | *pha-1 (e2123); him-5(e1490); otIs544 [cho-1(fosmid)::SL2::mCherry::H2B + pha-1(+)] V* |
| NC190 | *Dpy-20(e1282) IV; wdIs6 [del-1::GFP] II* |
| OH4128 | *evIs82b [unc-129::GFP + dpy-20(+)] IV; juIs76 [ unc-25::gfp + lin-15(+)] II* |
| | *ynIs37 [flp-13p::GFP] III* |
| KRA235 | *pha-1 (e2123); kasEx80 [poig-1_125bp::RFP#9 + pha-1 (+)]* |
| KRA236 | *pha-1 (e2123); kasEx81 [poig-1_125bp::RFP#10 + pha-1 (+)]* |
| KRA237 | *pha-1 (e2123); kasEx82 [poig-1_125bp::RFP#12 + pha-1 (+)]* |
| KRA252 | *pha-1 (e2123); kasEx91 [LIN-39 site mut poig-125bp::RFP#1 + pha-1 (+)]* |
| KRA253 | *pha-1 (e2123); kasEx92 [LIN-39 site mut poig-125bp::RFP#4 + pha-1 (+)]* |
| KRA254 | *pha-1 (e2123); kasEx93 [LIN-39 site mut poig-125bp::RFP#7 + pha-1 (+)]* |
| KRA256 | *pha-1 (e2123); kasIs2 [lin-39 $^{intron\ 1}$::tagRFP + pha-1 (+)]* |
| BC13337 | *dpy-5 (e907) I; sIs12928 [twk-46::GFP + pCeh361] V* |
| KRA22 | *pha-1 (e2123); kasEx22 [ilys-4::tagRFP + pha-1(+)]* |
| OH11954 | *lin-11::mCherry + myo-2::GFP V* |
| OH13105 | *him-5(e1490); otIs564 [unc-47(fosmid)::SL2::H2B::mChopti + pha-1(+)] V* |
| CZ13799 | *juIs76 [unc-25p::GFP + lin-15(+)] II* |
| CZ8332 | *juIs223 [ttr-39p::mCherry + ttx-3p::GFP] IV* |
| BC11799 | *dpy-5 (e907) I; sEx11799 [rCesF56E3.3::GFP + pCeh361]* |
| OP200 | *unc-119(ed3) III; wgIs200 [alr-1::TY1::EGFP::3xFLAG + unc-119(+)] X* |
| OP536 | *unc-119(tm4063) III; wgIs536 [irx-1::TY1::EGFP::3xFLAG + unc-119(+)] I* |
| OP395 | *unc-119(tm4063) III; wgIs395 [unc-30::TY1::EGFP::3xFLAG + unc-119(+)]* |
